# SWI/SNF coordinates transcriptional activation through Rpd3-mediated histone hypoacetylation during quiescence entry

**DOI:** 10.1101/426288

**Authors:** Marla M. Spain, Keean C.A. Braceros, Toshio Tsukiyama, Fred Hutchinson Cancer Research Center, Division of Basic Sciences

## Abstract

Whether or not a cell chooses to divide is a tightly regulated and extremely important decision. Cells from yeast to human are able to reversibly exit the cell cycle in response to environmental changes such as nutritional changes or removal of growth cues to become quiescent. An inappropriate response to environmental cues can result in overproliferation which can lead to cancer, or a failure to proliferate which can result in developmental defects, premature aging and defects in wound healing. While many of the cell signaling pathways involved in regulating cellular quiescence have been identified, how these pathways translate their messages into transcriptional outputs is not well characterized. We previously showed that the histone deacetylase Rpd3 mediates global histone deacetylation and transcription repression upon quiescence entry. How the activation of quiescence-specific genes occurs in the midst of this transcriptionally repressive environment is not well understood. We show that the SWI/SNF chromatin remodeling complex activates quiescence specific genes to promote entry into quiescence. We additionally show that SWI/SNF binding early during quiescence entry is important for facilitating localization of the transcriptional activator Gis1, as well as histone H4 hypoacetylation in coding regions later on. The increase in H4 acetylation that we observe at Snf2-regulated genes upon Snf2 depletion corresponds to a decrease in promoter-bound Rpd3, suggesting that Snf2 remodels chromatin not only to facilitate activator binding, but also the binding of Rpd3. These observations provide mechanistic insight as to how quiescence-specific genes can be activated in the face of global deacetylation and transcription repression.

## Introduction

Cells from yeast to human have the ability to voluntarily exit the cell cycle and become quiescent in response to environmental changes such as withdrawal of growth cues or nutritional depletion (Cheung and Rando, 2013; Coller et al., 2006; Valcourt et al., 2012; Werner-Washburne et al., 1993). Faithful regulation of cellular quiescence is essential for the health of all organisms, as its misregulation can lead to developmental defects, defects in wound healing, premature aging and cancer (Amrani et al., 2011; Bischoff, 1990; Cheung and Rando, 2013; Fausto, 2004; Rodgers et al., 2014; Schaniel et al., 2011). Quiescent cells across organisms share many characteristics such as exit from the cell cycle with G1 DNA content, more compact chromatin, decreased transcription and translation, decreased autophagy, decreased cell size, and increased stress resistance (De Virgilio, 2012; Dhawan and Laxman, 2015; Gray et al., 2004; Valcourt et al., 2012). Additionally, the main signaling pathways known to be involved in regulating quiescence entry and exit including the TORC, Ras/PKA, and AMPK pathways are conserved from yeast to human (De Virgilio and Loewith, 2006; Fabrizio et al., 2001; Longo and Fabrizio, 2012; Tatchell, 1986; Thevelein and de Winde, 1999). How these quiescence-specific signals are translated into gene regulatory responses, however, is still not well understood.

What we know about transcriptional regulation in response to nutrient limitation in budding yeast, is that it is complex, with a wide array of signaling pathways and transcription factors converging to facilitate successful passage through the diauxic shift (DS) where cells switch from fermenting glucose to utilizing ethanol, and entry into stationary phase and quiescence (DeRisi et al., 1997; Galdieri et al., 2010; Zhang et al., 2009). For example, the PKA and TOR pathways negatively regulate quiescence entry, whereas Rim15 and Snf1 promote it (Reviewed in (Galdieri et al., 2010)). Upon nutrient limitation, Rim15 responds to inhibition of the PKA and TOR pathways to facilitate binding of the stress response transcription factors Msn2/4 and the post-diauxic shift transcription factor Gis1 to the promoters of their target genes (Bontron et al., 2013; Pedruzzi et al., 2000; Swinnen et al., 2006; Zhang et al., 2009). These factors bind to specific DNA sequences termed STRE (stress-responsive elements) for Msn2/4 (Martínez-Pastor et al., 1996), or PDS (post-diauxic shift) elements in the case of Gis1 (Pedruzzi et al., 2000), to upregulate genes that are important for entry into stationary phase and quiescence. How the binding of these factors activates transcription, as well as what other factors are involved, however, is not known.

The positive correlation between histone acetylation and active transcription in vegetatively growing cells is well established. Cellular quiescence, however, is marked by decreased histone acetylation and transcription (Cheung and Rando, 2013; De Virgilio, 2012; McKnight et al., 2015). In fact, our lab previously showed that the histone deacetylase Rpd3 mediates global histone deacetylation to facilitate a dramatic genome-wide transcription shutdown during quiescence entry in budding yeast (McKnight et al., 2015). In this way, diauxic shift, post-diauxic shift and stationary phase genes must be upregulated in the midst of global histone deacetylation (McKnight et al., 2015), decreased global acetyl CoA levels (Cai et al., 2011; Shi and Tu, 2013), and a chromatin environment that is generally refractory to transcription (McKnight et al., 2015; Piñon, 1978; Rutledge et al., 2015). These genes, such as those required to transition from fermenting glucose to utilizing ethanol and later to metabolizing acetate (DeRisi et al., 1997; Galdieri et al., 2010; Gasch et al., 2000; Pedruzzi et al., 2000; Radonjic et al., 2005), or for fortifying the cell wall and increasing thermotolerance (Li et al., 2013; Shimoi et al., 1998), are likely activated by an alternative mechanism. We show that the catalytic subunit of the Swi/Snf chromatin remodeling complex, Snf2, regulates transcription of a subset of post-diauxic genes by promoting histone H4 hypoacetylation in coding regions. Depletion of Snf2 early during the quiescence entry process causes defects in binding of the transcriptional activator, Gis1, and the histone deacetylase Rpd3. Snf2-depletion additionally results in increased H4 acetylation over the coding regions of these genes. Based on these observations, we propose that Snf2 facilitates transcription of this class of genes at least partly through Rpd3-dependent H4 hypoacetylation. Overall, our results provide mechanistic insight as to how post-diauxic shift genes are specifically activated in the midst of a repressive chromatin landscape.

## Results

### Snf2 is required for Q entry

Dynamic changes in both transcription and chromatin structure occur during quiescence (Q) entry (Aragon et al., 2008; McKnight et al., 2015; Piñon, 1978; Rutledge et al., 2015). We hypothesized that chromatin remodeling might be important for regulating these changes. To test this hypothesis, we screened a number of chromatin remodeling mutants in an auxotrophic background for defects in Q entry, and discovered that the SWI/SNF complex was the strongest hit. In support of these results, the Snf6 subunit of the SWI/SNF complex was shown to be required for Q cell formation (Li et al., 2015), and Snf2 was predicted to be a quiescence specific transcription factor by computational analyses (Reimand et al., 2012). Additionally, the SWI/SNF complex is a well-established regulator of transcription, known for activating glucose-repressed genes (Abrams et al., 1986) and genes involved in metabolic regulation and/or stress through its nucleosome remodeling activity (Awad et al., 2017; Côté et al., 1994; Dutta et al., 2014; Erkina et al., 2008; Geng and Laurent, 2004; Peterson et al., 1994; Qiu et al., 2016; Shivaswamy and Iyer, 2008; Wang et al., 2018).

To confirm that the SWI/SNF complex is important for Q entry, we grew prototrophic cells harboring a deletion of the catalytic subunit, Snf2, for seven days, purified them on a Percoll density gradient (Allen et al., 2006) and assessed Q cell yield. Compared to the wild type cells, the *snf2Δ* cells grew extremely slowly, flocculated heavily and produced virtually no Q cells (Figure 1A, B). This result indicates that the SWI/SNF complex is indeed involved in regulating key events that are required for Q entry. Quiescence entry is a lengthy process, beginning when the glucose in the media starts to wane, and ending nearly a week later (Allen et al., 2006; Lillie and Pringle, 1980). When about half of the glucose has been consumed from the media, wild type budding yeast cells gradually stop cycling and arrest in G1 (Allen et al., 2006; François and Parrou, 2001; Li et al., 2013; Lillie and Pringle, 1980; Parrou et al., 1999). The last S-phase is nearly complete in the population by the DS (Li et al., 2013), and G1 arrest is complete by about 20 hours after the DS ((Li et al., 2013); Figure S1A). Additionally, quiescent cell yield has recently been correlated with the length of time cells spend in G1 during logarithmic growth (Miles and Breeden, 2016, 2017), suggesting that the timing of the G1 arrest with respect to the DS is important for Q entry. To rule out the possibility that abnormal cell-cycle progression is responsible for the Q entry defect observed in *snf2Δ* cells, we analyzed cell-cycle profiles of wild type and *snf2Δ* cells during Q entry by fluorescence activated cell sorting (FACS). The *snf2Δ* cells arrested in G1 at around the same time relative to the DS as the wild type cells, despite growing twice as slowly (Figure S1A). These results indicate that the Q entry defects observed in *snf2Δ* cells are not simply due to a failure to arrest in G1 appropriately.

**Figure 1.**
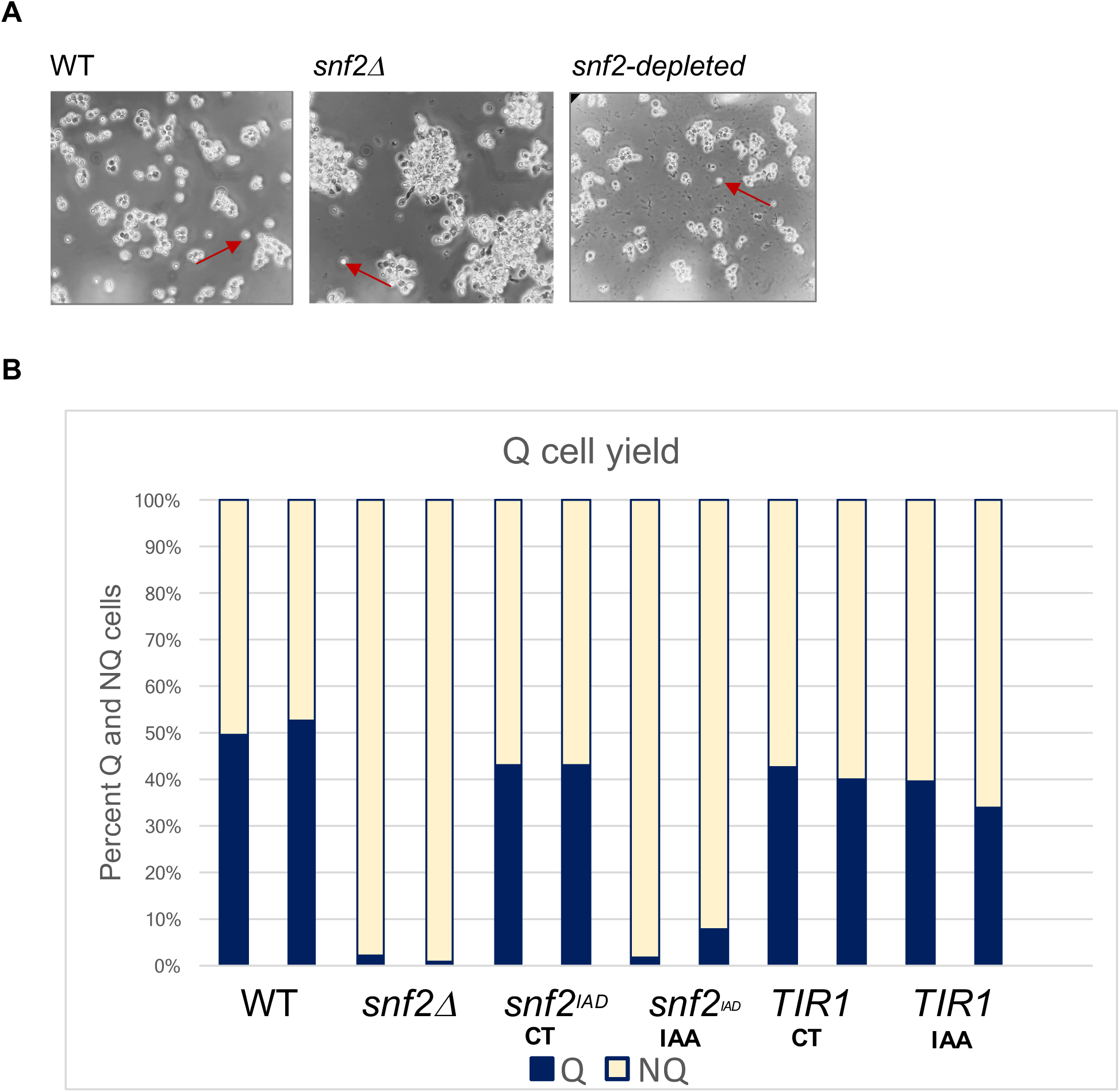
Snf2 is required for Q cell formation. **A) (top)** Images of WT and *snf2*Δ cells taken 3 days after inoculation. The small shiny round cells in the WT image, indicated by the red arrows, are Q cells. *snf2*Δ cells are flocculated and virtually no Q cells are visible. **(bottom)** Q cell yield measured at Day 7 indicates that *snf2*Δ cells form virtually no Q cells. **B) (top)** Western blot with anti-HSV antibody recognizing the Snf2 protein in untreated cells (0’) or in IAA-treated cells 5’-60’ after the addition of IAA. Antibody to H2B was used as a loading control. **(bottom)** Images of WT and Snf2-depleted cells taken 3 days after inoculation. Very few Q cells are present in the Snf2-depleted sample. **C)** Percentage of Q and NQ (cells that did not enter Q after 7 day culture) cells purified 7 days after inoculation in WT and *snf2*Δ cells; *snf2*-*AID* cells untreated and treated with IAA in log phase; and cells possessing the ubiquitin ligase complex but not the auxin tag on Snf2 untreated or treated with IAA in log phase.

### The requirement for Snf2 during Q entry is strongest just before the DS

As mentioned above, the process of Q entry begins even before glucose has been depleted from the media, and lasts nearly a week (Allen et al., 2006; Li et al., 2015). Snf2 could be involved in regulating any or even multiple stages of the process. Because the null strain precluded us from examining specific temporal contributions of Snf2 to the quiescence entry process, we generated a degron strain in which the Snf2 protein could be rapidly degraded by the addition of the plant hormone auxin to the media (Nishimura et al., 2009). Degradation of Snf2 began in as early as 5 minutes after auxin addition, and was fairly complete shortly thereafter (Figure S1B), allowing us to control Snf2 protein levels temporally. To compare the effects of Snf2 depletion on Q entry with those of *snf2Δ*, we added auxin (IAA) to exponentially growing (log) cells and allowed them to grow for seven days. Similar to *snf2Δ*, the *snf2*-*AID* cells produced almost no Q cells (Figure 1B). Additionally, Snf2-depleted cells exhibited substantial cell death beginning at around two days post DS (Figure 1A). Cells expressing a functional E3 ubiquitin ligase complex (*TIR1*) but not the *snf2*-*AID* degron produced nearly wild type levels of Q cells (Figure 1B), and showed no signs of cell death. These results indicate that the phenotypes observed in *snf2*-*AID* cells were due to depletion of Snf2 (Figure 1B).

Having established a system to control Snf2 protein levels temporally, we depleted Snf2 at various time points during the Q entry process to more closely determine when Snf2 functions to promote quiescence. Depleting Snf2 in log cells resulted in the greatest decrease in Q cell yield, whereas depleting Snf2 at or shortly after the DS produced far higher levels of Q cells (Figure 2A; S2A). Q cell yield decreased and cell death increased with increasing time of depletion prior to the DS (Figure S2A), suggesting that the contribution of Snf2 to Q entry begins during log phase. To determine whether the observed defects reflected a specific function of Snf2 just prior to the DS, rather than cumulative effects from its role in log cells, we depleted Snf2 specifically for three hours during late log phase or within the last four hours before the DS. To do so, we washed out the IAA after the depletion period by filtering the cells from the media and re-suspending them in spent media from a duplicate culture. We then allowed the cells to enter Q. Washing out the IAA at DS-3h after a 3-hour depletion resulted in a substantial rescue of Q cell yield (Figure 2B; Figure S2C). The Snf2 protein was detectable by western blot by about 30 minutes after the IAA was washed out, and FACS profiles of the rescued cells were identical to the control cells (Figure 2B; Figure S2B). To the contrary, when we washed out the IAA after depleting Snf2 from about five hours prior to the DS through the DS, we were unable to rescue Q cell yield (Figure 2C; Figure S2D). The lack of rescue, along with the finding that depleting Snf2 after the DS has little to no effect on Q cell yield, indicates both that Snf2 performs an essential function during the last several hours prior to the DS, and that Snf2 expression and/or protein levels are regulated during this transition. Consistent with this possibility, Snf2 expression measured by nascent RNA sequencing (RNA-seq) is greatly reduced at DS+24h as compared to exponentially growing cells (Figure S2E). Additionally, protein levels for Snf2 peak just prior to the DS and decrease afterward (Figure S2F). Overall, these results point to a specific vital function for Snf2 in the hours preceding the DS.

**Figure 2.**
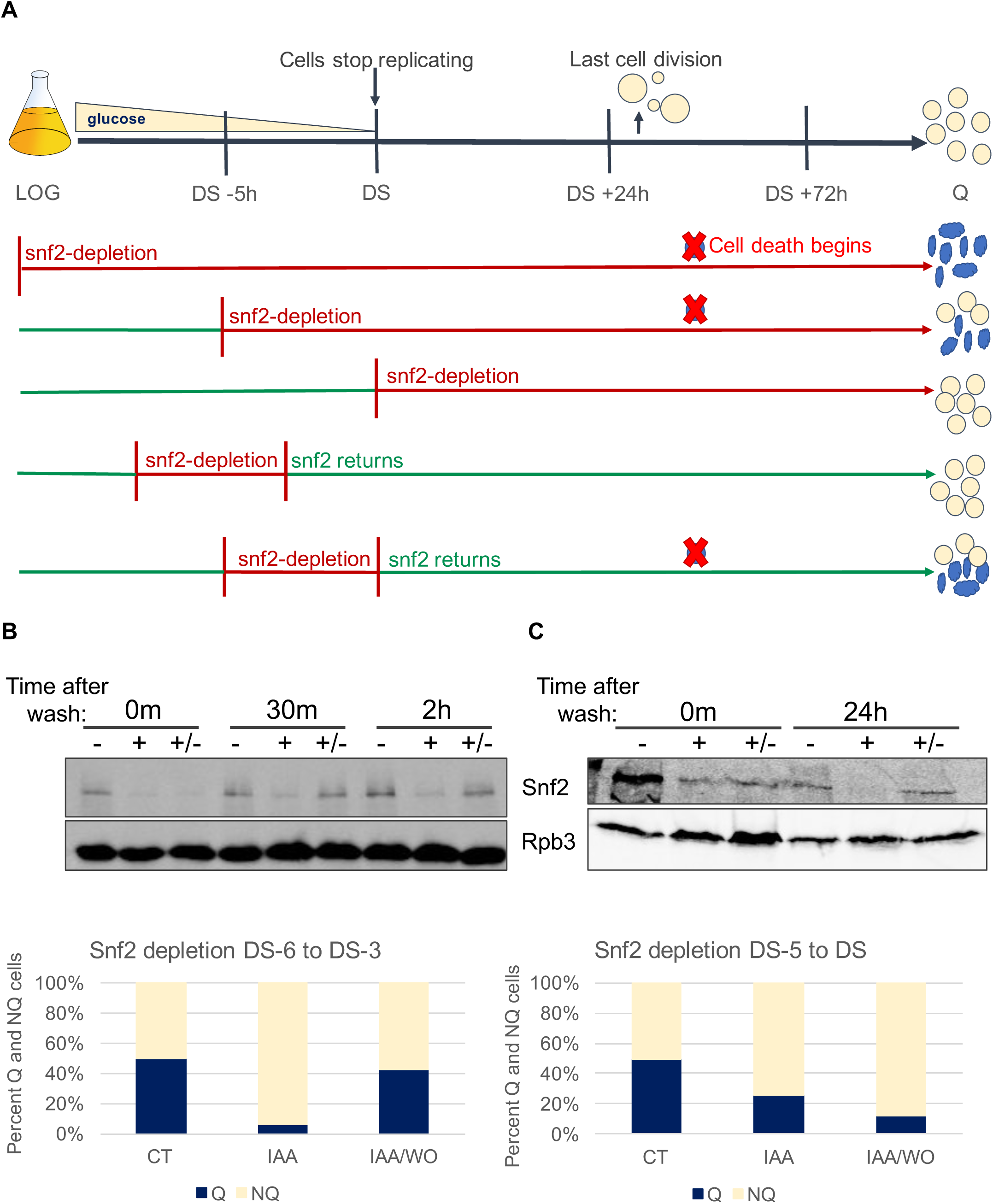
The requirement for Snf2 during Q entry is strongest just prior to the DS. **A)** Schematic drawing of the Q entry process and summary of auxin depletion experiments indicating that Snf2 is most important prior to the DS. On the right, yellow and blue cells denote live and dead cells, respectively. **B) (top)** Western blot with antibody to HSV-tagged Snf2 and Rpb3 as a control. Untreated (-), treated (+) or treated and washed (+/-) samples were harvested at the indicated times after washout. Treated cells were grown in the presence of auxin from DS-6h to DS-3h. **(bottom)** Percent Q and NQ cells produced by untreated, treated or treated and washed cells harvested 7 days after inoculation. **C)** The same as in B except that cells were treated with auxin from DS-5h to the DS.

### Snf2 regulates transcription of genes required for the diauxic shift

Having established that Snf2 is particularly important prior to the DS, we sought to understand how it might function during this time period. To this end, we measured Snf2 occupancy genome-wide in log cells and at various time points throughout the Q entry process by chromatin immunoprecipitation followed by deep sequencing (ChIP-seq). Genome-wide K-means clustering analysis revealed that the global binding pattern for Snf2 is dynamic from log to DS-2h, but is fairly stable thereafter (Figure 3A). In log cells, Snf2 bound on either side of the nucleosome depleted region (NDR) of genes involved mainly in cytoplasmic translation and ribosome biogenesis, in agreement with previous studies (Dutta et al., 2014; Yen et al., 2012) (Figure 3A). The Snf2 binding profile two hours prior to the DS (DS-2h) was very different from the log profile (Figure 3A), consistent with our observation that Snf2 is specifically required in the last few hours prior to the DS (Figure 2A). The genes bound at this time point were mainly involved in carbohydrate metabolism and glycolysis, indicating that Snf2 may play a role in regulating these metabolic processes. While the binding patterns were not identical throughout the course of the Q entry process, the three clusters of genes that were most highly occupied by Snf2 at DS-2h (clusters 1-3) remained highly bound from the DS through Q (Figure 3A). Overall, the similarity in the Snf2 binding pattern from DS-2h through Q suggests that the binding pattern of SWI/SNF at its target genes is globally established in the waning hours before the DS.

**Figure 3.**
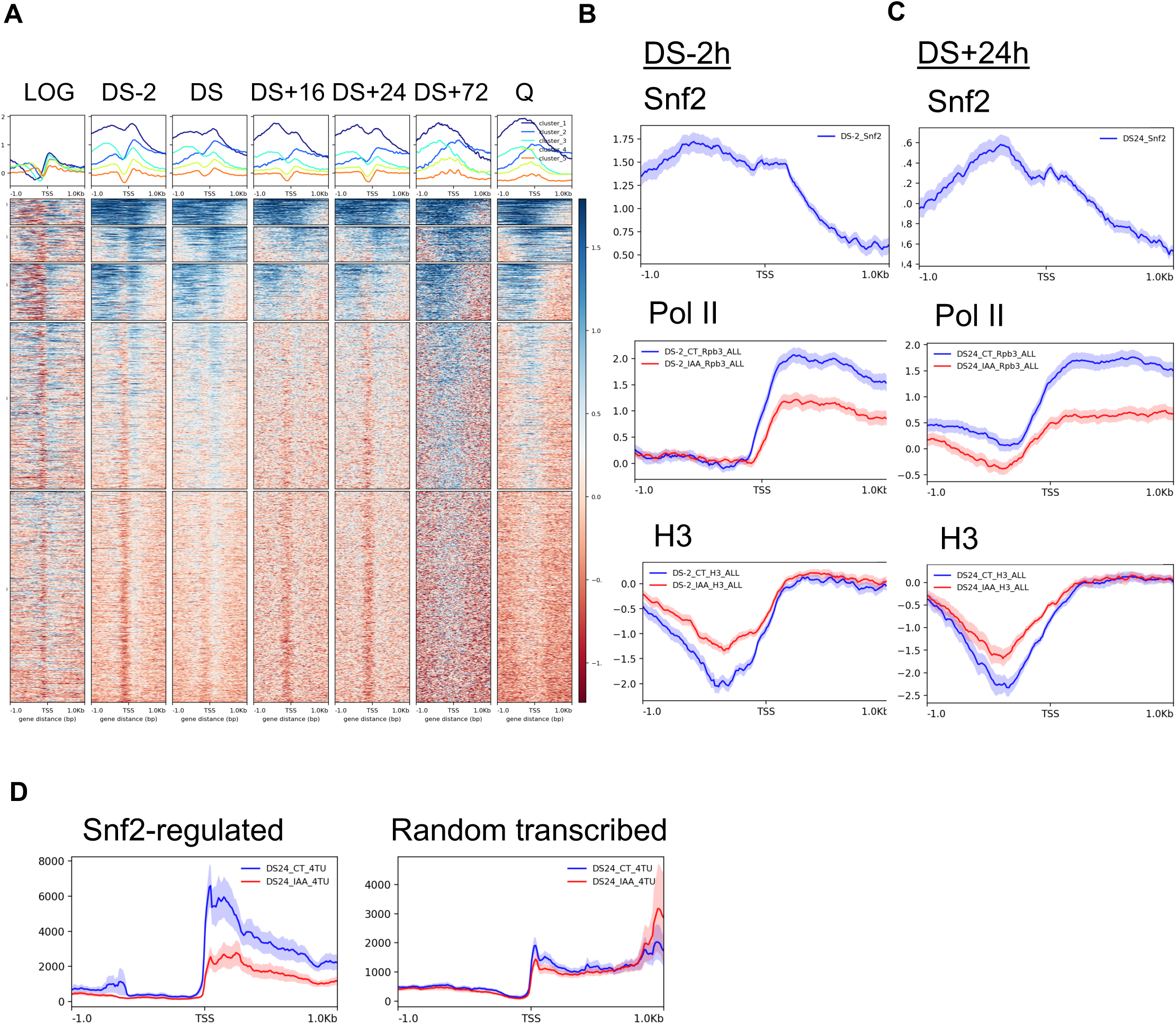
Snf2 binds and regulates genes required for passage through the diauxic shift. **A)** Heatmap showing K-means clustered Snf2-FLAG ChIP-seq data at the indicated times. Each column shows -/+ 1kb on either side of the transcriptional start site. **B)** Meta-analyses showing Snf2 (top), Pol II (Rpb3) (middle) and H3 (bottom) occupancy at genes bound and regulated by Snf2 at DS-2h. Shaded lines represent the standard error of the mean (SEM) for all meta-plots. **C)** The same as in B, except for genes bound by Snf2 at DS+24h. **D)** Meta-analysis of 4TU RNA-seq data at DS+24 in CT and auxin-treated cells at the genes bound and regulated by Snf2 (left) and at a set of randomly chosen transcribed genes (right).

To determine whether Snf2 regulates transcription of the genes it targets, we analyzed RNA polymerase II (Pol II) occupancy at two time points (DS-2 and DS+24 hours) by ChIP-seq of the Rpb3 subunit in wild type and Snf2-depleted cells. We added IAA to deplete Snf2 four hours prior to the DS for all of the following experiments. We chose the DS-2h timepoint because it is within the critical window for Snf2 activity. The DS+24h timepoint occurs after the cells are completely arrested in G1, but before the Snf2-depleted cells begin to die.

Of the 700 Snf2-bound genes at DS-2h (defined as genes exhibiting greater than log_2_ 0.75 Snf2 occupancy), 60 exhibited decreased Pol II occupancy in Snf2-depleted cells at DS-2h (Figure 3B, top). These genes are involved in glycolysis, gluconeogenesis and cell wall fortification. Concurrent with a decrease in Pol II occupancy (Figure 3B, middle), increased histone occupancy over the nucleosome depleted regions (NDRs) of these genes, measured by histone H3 ChIP-seq (Figure 3B, bottom), suggests that Snf2 remodels chromatin at their promoters to promote transcription. As a control, we analyzed Pol II and H3 occupancy of 96 randomly chosen transcribed genes with Snf2 binding below our chosen threshold, and observed negligible effects of Snf2 depletion on Pol II and H3 occupancy at this set of genes in Snf2-depleted cells (Figure S3A). Around the time that we observe Snf2 binding to genes involved in glycolysis and gluconeogenesis, yeast cells approaching the DS reduce glycolytic flux and increase the production of storage compounds such as glycogen and trehalose (François and Parrou, 2001; Lillie and Pringle, 1980; Parrou et al., 1999; Zampar et al., 2013). After the DS, the cells switch to gluconeogenesis and aerobic respiration to consume the ethanol produced during fermentation (François and Parrou, 2001; Zampar et al., 2013). Targeting and regulation of genes involved in both glycolysis and gluconeogenesis just prior to the DS indicates that Snf2 is involved in regulating transcription of genes required for the DS.

### Snf2 promotes transcription of genes involved in cell wall fortification

Alterations in the yeast cell wall that occur on the way to stationary phase mediate the altered light scattering, increased stress resistance and thermotolerance, characteristic of quiescent cells (Li et al., 2015; Li et al., 2013; Shimoi et al., 1998). Genes that mediate these effects, such as *SED1*, *ECM33* and *CWP2*, are essential for quiescence entry (Li et al., 2015; Li et al., 2013). These genes are also required for the altered light scattering observed in quiescent cells. We observe that Snf2 binds and regulates these and other genes involved in cell wall organization at DS-2h. Additionally, FACS analysis of WT and Snf2-depleted cells at the DS and DS+24h indicate that Snf2-depleted cells do not adopt the shift in light scattering mediated by *ECM33*, *SED1* and *CWP2* in Q cells (compare blue to orange at DS+24h)((Li et al., 2013); Figure S2B). Together, these results suggest that in addition to regulating glycolysis and gluconeogenesis, Snf2 promotes Q entry through facilitating transcription of genes required for cell wall fortification.

### Snf2 regulates carbohydrate metabolism at multiple stages of quiescence entry

At DS+24h, 67 Snf2-bound genes (Figure 3C, top), exhibited decreased Pol II occupancy in coding regions and increased histone occupancy over the NDR upon Snf2 depletion (Figure 3C, middle, bottom). Gene ontology analysis revealed that the Snf2-bound genes are involved in carbohydrate metabolism and pyridine nucleotide metabolism. No changes in Pol II or H3 occupancy were observed at either 73 randomly chosen transcribed genes (with Snf2 < log_2_ 0.75) (Figure S3B, top) or in a strain with the *TIR1* construct but no Snf2-degron at DS+24h (Figure S3B, bottom), indicating that the observed effects are specific to Snf2-depletion. Additionally, no enrichment over input was observed at either the 73 random genes or the 67 Snf2-bound genes in ChIP-seq samples with no FLAG antibody (no primary antibody control)(Figure S3C). Overall, these results suggest that Snf2 regulates metabolism not only during the diauxic shift, but at later stages in the Q entry process as well.

To ensure that Pol II ChIP-seq was a faithful indicator of active transcription, we performed nascent RNA-seq (4TU-seq) in wild type and Snf2-depleted cells in log phase and at DS+24h. Cells grown to log phase or to DS+24 were incubated with 4-thiouracil for 6 minutes to label nascent RNA (Duffy et al., 2015; Duffy and Simon, 2016; Warfield et al., 2017) and immediately spun down and flash frozen. Labeled RNA was extracted, biotinylated, and purified using streptavidin beads as described previously (Duffy et al., 2015; Duffy and Simon, 2016; Warfield et al., 2017). For unknown reasons, a substantial amount of unlabeled RNA bound to the streptavidin beads at DS-2h, preventing us from utilizing this technique at this time point. Overall, there was good agreement between the 4-TU RNA-seq replicates, and between the 4-TU RNA-seq and Pol II Chip-seq data in both log and DS+24h cells (Figure S3D), demonstrating that Pol II ChIP-seq is an appropriate way to measure the rate of active transcription. In support of this conclusion, genes exhibiting a decrease in Pol II at DS+24h in Snf2-depleted cells also showed a decrease in 4TU signal at that time, whereas the 73 randomly chosen control genes did not (Figure 3D).

### Snf2 regulates chromatin structure and transcription at post-diauxic genes

To investigate the mechanism by which Snf2 might regulate transcription of Q-specific genes in the last few hours before the DS, we analyzed Pol II binding at DS-2h and DS+24h specifically at the 700 genes bound by Snf2 at DS-2h. Because different sets of genes are activated at various time points during Q entry, we analyzed the temporal transcriptional profiles of genes bound by Snf2 at DS-2h to identify the crucial regulatory events occurring at this time point. K-means clustering analysis revealed four distinct clusters of transcribed genes (Figure S4A). We were surprised to observe what appeared to be two distinct relationships between Snf2 binding and transcription. At the genes in Clusters 1 and 4, Snf2 binding and transcription are concurrent (Figure S4A). In contrast, Snf2 binding precedes transcription at the genes in Cluster 2 (Figure S4A). ChIP-seq of Snf2 and Pol II in log cells, and at DS-2h, DS, DS+16h and DS+24h, supports these observations as Cluster 1 and 4 genes are bound by Snf2 at DS-2h and are transcribed at that time point (Figure S4B), whereas the majority of the genes in Cluster 2, although bound by Snf2 at DS-2h, are not activated until two hours later at the DS (Figure S4C).

To further investigate the mechanism by which Snf2 promotes Q entry, we sought to determine whether and how Snf2 might be regulating these distinct sets of genes. To this end, we went back to the 700 genes bound by Snf2 at DS-2h, and defined two distinct subclasses of Snf2-target genes in the following manner, defining “active transcription” as Pol II occupancy of greater than log_2_ 0.25. From all of the genes that were bound by Snf2 at DS-2h, we took the genes that are actively transcribed at that time. From this group, we pulled out Snf2-regulated genes (with more than a log_2_ 0.5 decrease in Pol II occupancy in Snf2-depleted cells). We defined this group as Class I (60 genes). From the Snf2-bound genes that are not transcribed at DS-2h, we took genes that are transcribed at DS+24. From this group, we identified Snf2-regulated genes (with a greater than log_2_ 0.5 decrease in Pol II occupancy in Snf2-depleted cells). This group was defined as the Class II genes (41 genes).

Both classes of genes, in addition to by definition being bound by Snf2 at DS-2h, were robustly bound by Snf2 at DS+24h (Figure 4A and B, top, 4C, D). Consistent with the definition, the Class I genes exhibited a decrease in Pol II occupancy at DS-2h (Figure 4A, middle-left, 4C) and an increase in H3 occupancy over the NDR (Figure 4A, bottom-left, 4C) upon Snf2 depletion, suggesting that Snf2 remodels chromatin to promote transcription of this set of genes at DS-2h. Unexpectedly, both Pol II and H3 occupancy at Class I genes recover to near normal levels at DS+24h (Figure 4A, middle- and bottom-right, 4C), suggesting that the requirement of Snf2 for these genes is transient. The Class II genes, despite being strongly bound by Snf2 at DS-2h, are not transcribed at this time point (Figure 4B, middle-left, 4D). They do, however, show a slight increase in histone H3 occupancy over the NDR at DS-2h in Snf2-depleted cells (Figure 4B, bottom-left, 4D), supporting a model that Snf2 remodels chromatin at this time to facilitate later transcription activation (Figure 4E). In further support of this hypothesis, the substantial decrease in Pol II occupancy at Class II genes at DS+24h upon Snf2 depletion (Figure 4B, middle-right) is accompanied by an equally large increase in H3 occupancy over the NDR (Figure 4B, bottom-right).

**Figure 4.**
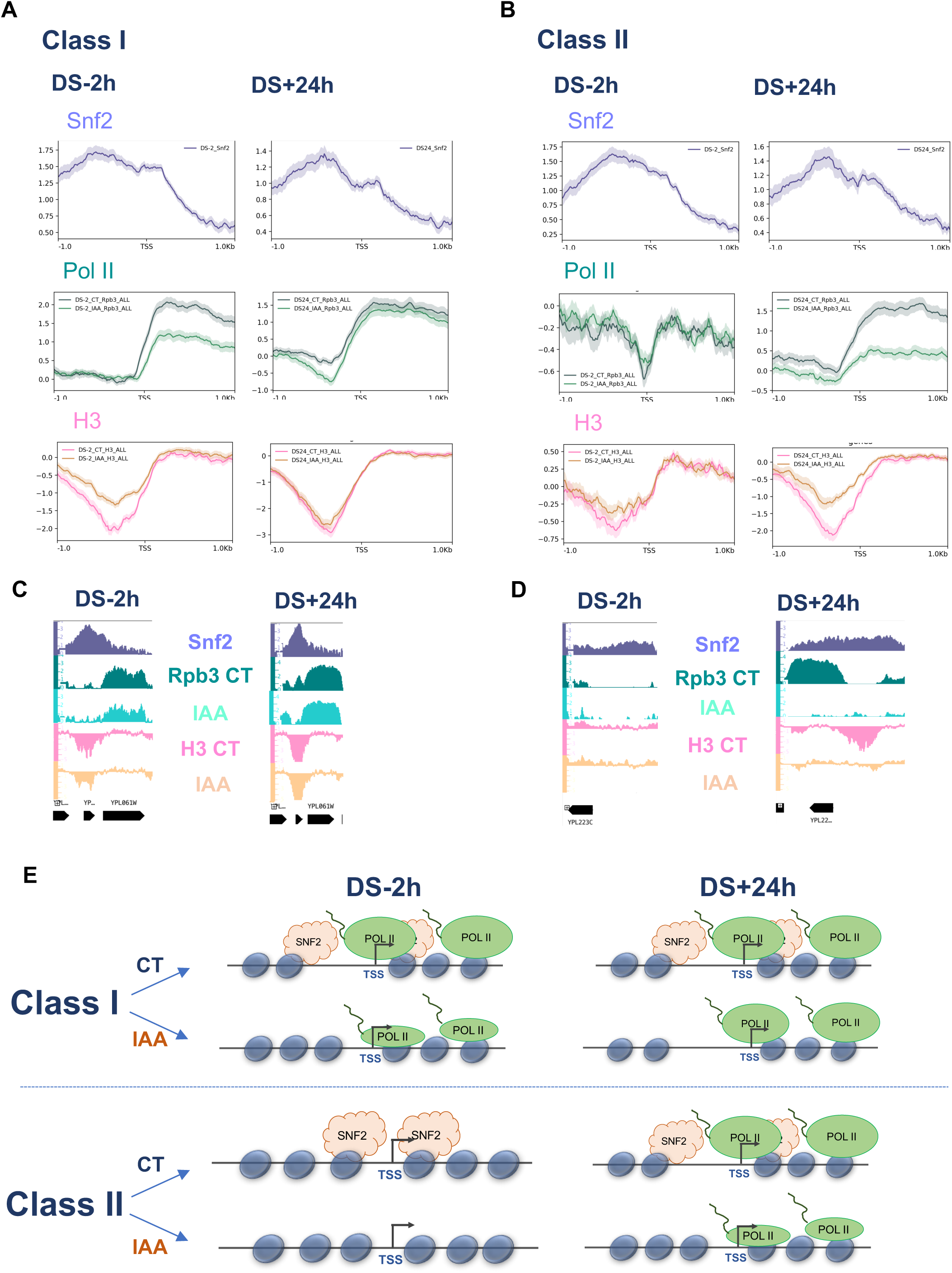
Snf2 regulates transcription and chromatin structure at post-diauxic shift genes. **A)** Meta-analyses showing Snf2 binding (top), Pol II occupancy (middle) and histone H3 occupancy (bottom) in CT and Snf2-depleted cells at DS-2h (left) and DS+24h (right) at Class I genes. **B)** The same for Class II genes. **C)** Screen shot of a representative Class I gene at DS+2 and DS+24 showing Snf2 binding (top), Pol II occupancy (middle), and H3 occupancy (bottom) in CT and Snf2-depleted (IAA) cells. **D)** The same for a representative Class II gene. **E)** Schematic drawing of the transcription and chromatin effects observed at Class I and Class II genes in Snf2-depleted cells.

### Snf2 facilitates activator binding at post-diauxic shift genes

The small increase in H3 occupancy over the NDR of Class II genes at DS-2h upon Snf2 depletion led us to wonder whether Snf2 might facilitate activator binding at this set of genes. While Snf2 has most commonly been shown to be recruited in a gene-specific manner by transcriptional activators (Cosma et al., 1999; Yoshinaga et al., 1992; Yudkovsky et al., 1999) to remodel chromatin and promote transcription (Ryan et al., 1998), it is also known to facilitate activator binding at various genes (Burns and Peterson, 1997; Côté et al., 1994; Imbalzano et al., 1994; Kwon et al., 1994). To test our hypothesis, we searched for transcription factors known to regulate the Class II genes using YEASTRACT (Abdulrehman et al., 2011; Monteiro et al., 2008; Teixeira et al., 2006; Teixeira et al., 2014; Teixeira et al., 2018; Teixeira et al., 2016), and discovered that the post-diauxic transcription factor, Gis1, was one of the top hits. Gis1 is phosphorylated upon nutrient depletion, which promotes its recruitment to metabolic genes whose transcription is activated during Q entry (Bontron et al., 2013; Pedruzzi et al., 2000). At a global level, ChIP-seq of FLAG-tagged Gis1 at DS-2h and DS+24h revealed that Gis1 and Snf2 co-localized at many genes at DS+24h (Figure 5A). Additionally, Gis1 binding is reduced upon Snf2 depletion at Class II genes which also exhibit a robust decrease in transcription in Snf2-depleted cells (Figure 5B). Snf2 binding, however, was unaffected in *gis1Δ* (Figure S5A). Interestingly, while Gis1 also binds to Class I genes, neither its binding nor transcription is affected at these genes at this timepoint by Snf2 depletion (Figure 5B). These results, as well as the increased H3 occupancy over the NDR of Class II genes in Snf2-depleted cells, suggest that Snf2 remodels chromatin at the promoters of Class II genes to facilitate Gis1 binding and transcriptional activation.

**Figure 5.**
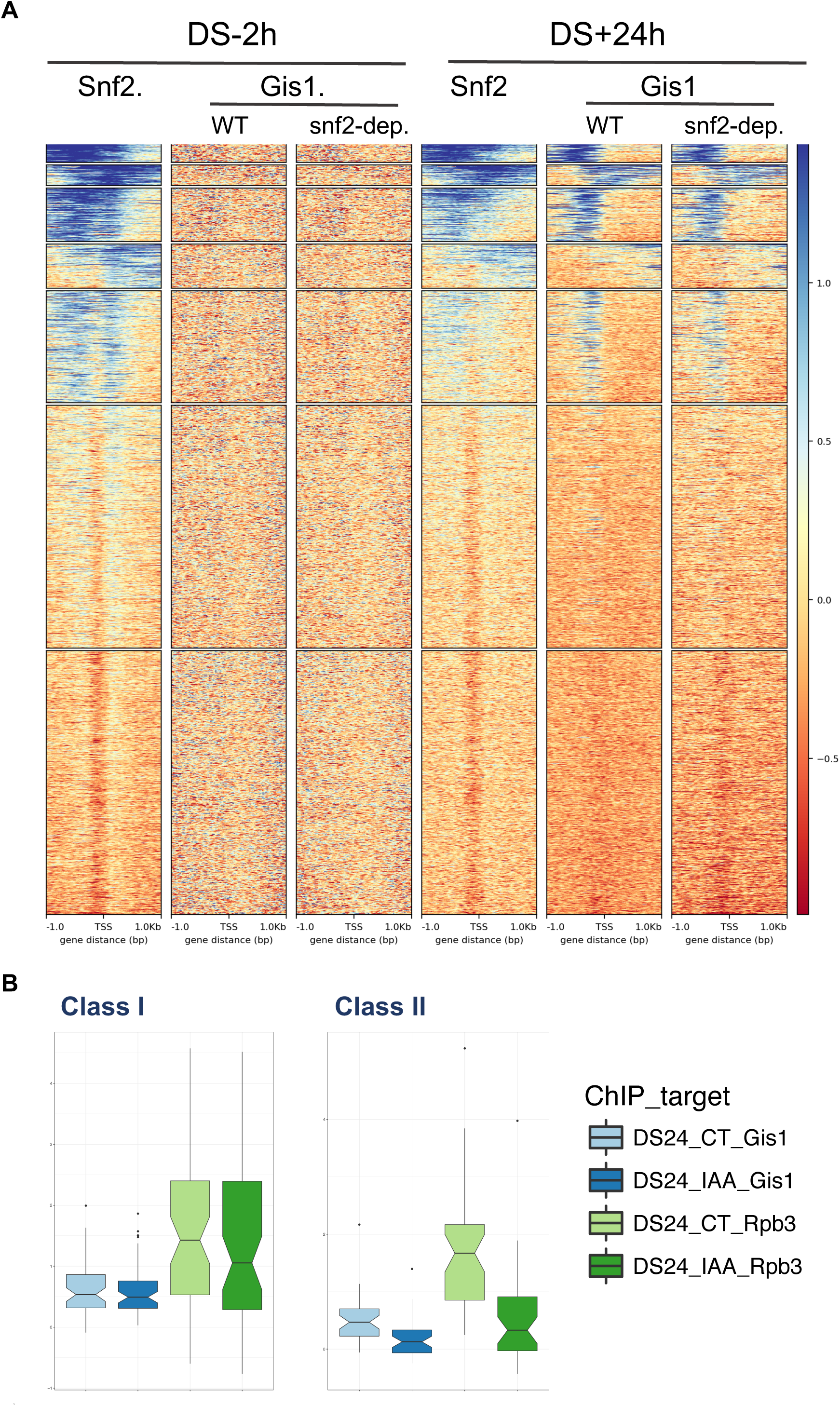
Snf2 facilitates activator binding at post-diauxic shift genes. **A)** Heatmap of K-means clustered Snf2 and Gis1 ChIP-seq data at DS-2h and DS+24h showing that the two factors co-localize at many genes at DS+24h. **B)** Box plots showing average Gis1 and Pol II binding (Rpb3) over the promoter and coding regions (Gis1) or the coding regions (Rpb3) at DS+24h at Class I (left) and Class II (right) genes.

### Snf2 promotes histone hypoacetylation in the coding regions of post-diauxic shift genes

Snf2 has a well-established relationship with histone acetylation, particularly with that mediated by the histone acetyltransferase Gcn5 (SAGA). For example, histone acetylation by SAGA is required for SWI/SNF binding to the *HO* (Mitra et al., 2006), *PHO8* (Reinke et al., 2001), and *CLN3* (Shi and Tu, 2013) promoters. Additionally, SWI/SNF specifically displaces nucleosomes acetylated by SAGA (Chandy et al., 2006; Hassan et al., 2006; Hassan et al., 2002). Histone acetylation also regulates the chromatin remodeling activity of SWI/SNF at stress response genes such as those in the Slt2 MAPK pathway (Sanz et al., 2012; Sanz et al., 2016), the heat shock response (Shivaswamy and Iyer, 2008), and the response to low glucose (Dutta et al., 2014).

To better understand the mechanism by which Snf2 binds and regulates transcription of post-diauxic shift genes, we measured histone acetylation by ChIP-seq genome-wide at DS-2h and DS+24h in wild type and Snf2-depleted cells with antibodies recognizing acetylation on H3 lysine 14 (H3K14ac) and collectively on H4 lysines 5, 8,12 and 16 (H4ac). Of the 426 genes that are induced at the diauxic shift and expressed at DS+24h in our dataset, 155 are bound by Snf2, including the 41 Class II genes. We compared H4 acetylation and transcription upon Snf2 depletion at the Class II genes with that of the 271 unbound induced genes. Unexpectedly, H4 acetylation levels decrease specifically in the coding region from DS-2h to DS+24h at the Class II genes as they are activated (Figure 6A), along with a much smaller decrease in total H3 occupancy across the coding region (Figure 6 C left, S6B). Upon Snf2 depletion, however, H4ac levels in the coding regions of these genes are increased back to the level of DS-2h (Figure 6A), whereas total H3 levels across the coding regions remain unchanged (Figure 6C left). These observations demonstrate that Snf2 plays a role in promoting H4 deacetylation in the coding regions of Class II genes, and further suggests that H4 hypoacetylation is required for activation of these genes. H3K14ac level was unaffected at Class II genes upon Snf2-depletion (Figure S6A), showing that the link between acetylation and transcription is unique to H4ac. At the Snf2-unbound induced genes, H4ac increases slightly from DS-2h to DS+24h, and neither H4ac nor transcription is affected upon Snf2 depletion at these genes (Figure 6A, C right). These results indicate that Snf2 may play roles in creating the anti-correlation between H4ac in coding regions and transcriptional activation.

**Figure 6.**
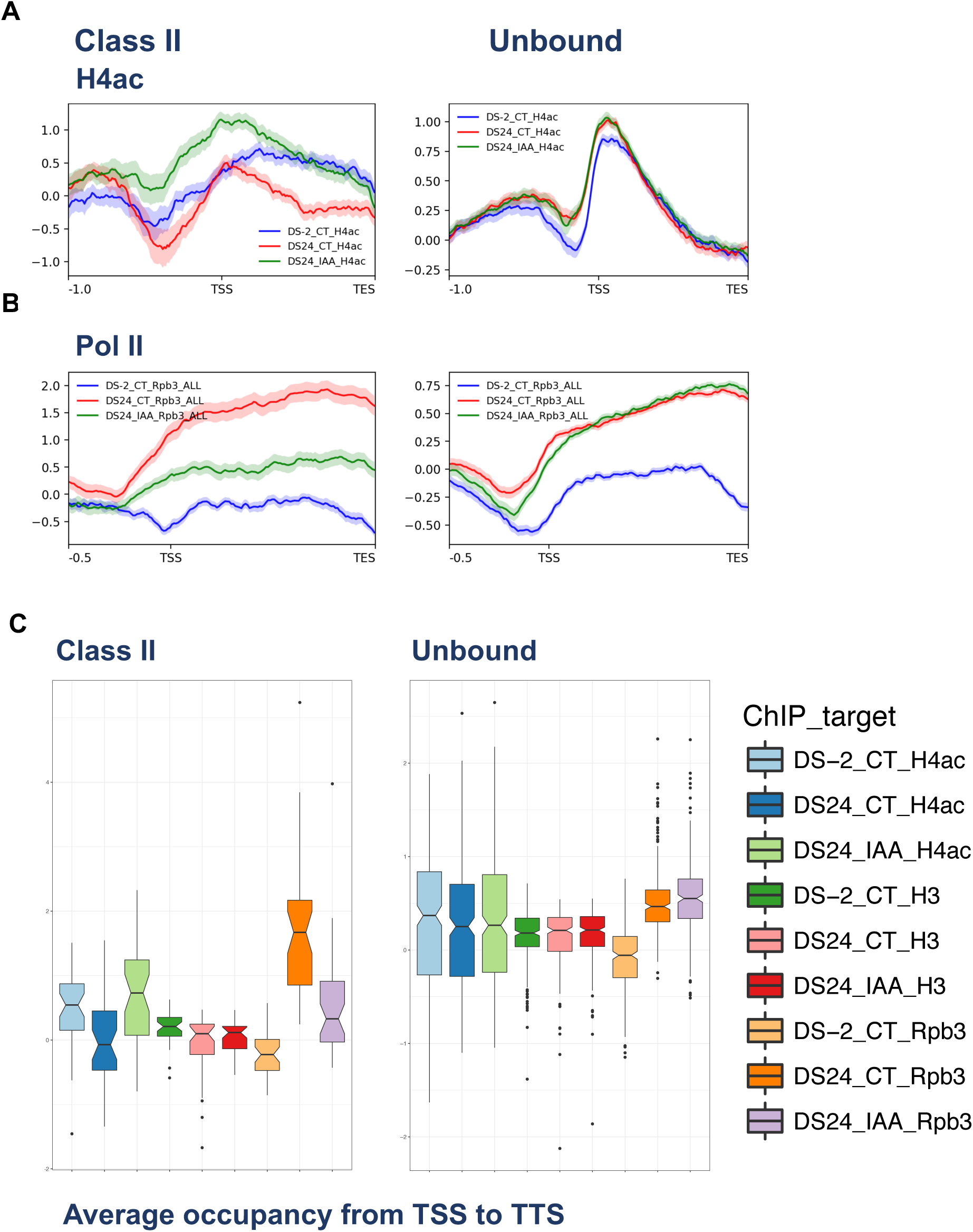
Snf2 promotes histone hypoacetylation in the coding regions of post-diauxic shift genes. **A)** Meta-analyses of H4ac ChIP-seq data at DS-2h (blue) and DS+24h (red) in CT cells, and at DS+24h in Snf2-depleted cells (green) at Class II and induced unbound genes. **B)** The same for Pol II ChIP-seq data. **C)** Box plots showing the average binding of H4ac, H3 and Pol II (Rpb3) across the coding regions of Class II (left) and Snf2-unbound genes that are also transcriptionally induced after the DS (right).

### Snf2 mediates Rpd3 binding to facilitate H4 hypoacetylation and transcription

To investigate how Snf2 might facilitate histone H4 hypoacetylation in the coding regions of post-diauxic shift genes, we measured genome-wide binding of the histone deacetylase Rpd3. While Rpd3 has been shown to mediate the global histone deacetylation and transcription repression that occurs during Q entry (McKnight et al., 2015), it is also known to deacetylate histones in the coding regions of active genes (Drouin et al., 2010; Govind et al., 2010; Keogh et al., 2005; Li et al., 2007). Strikingly, genome-wide K-means cluster analysis of Rpd3 ChIP-seq, along with Snf2, Pol II and H4ac, revealed that the genes most strongly bound by Rpd3 at DS+24h are also bound by Snf2, and are highly transcribed and hypoacetylated for H4 (Figure 7A, Cluster 2). This result demonstrates that the tight link between Snf2 targeting, H4 hypoacetylation in coding regions and active transcription can be identified through unbiased genome-wide analyses at DS+24h. At Class II genes, Rpd3 binds to the promoters, and its binding is strongly Snf2-dependent (Figure 7B). Together, these results suggest a model in which Snf2 remodels chromatin at the promoters of Class II genes prior to the DS (Figure 7C, top) to facilitate binding of Gis1 activator and Rpd3 afterward (Figure 7C, bottom). Histone H4 hypoacetylation mediated by this series of events then facilitates active transcription (Figure 7C, bottom).

**Figure 7.**
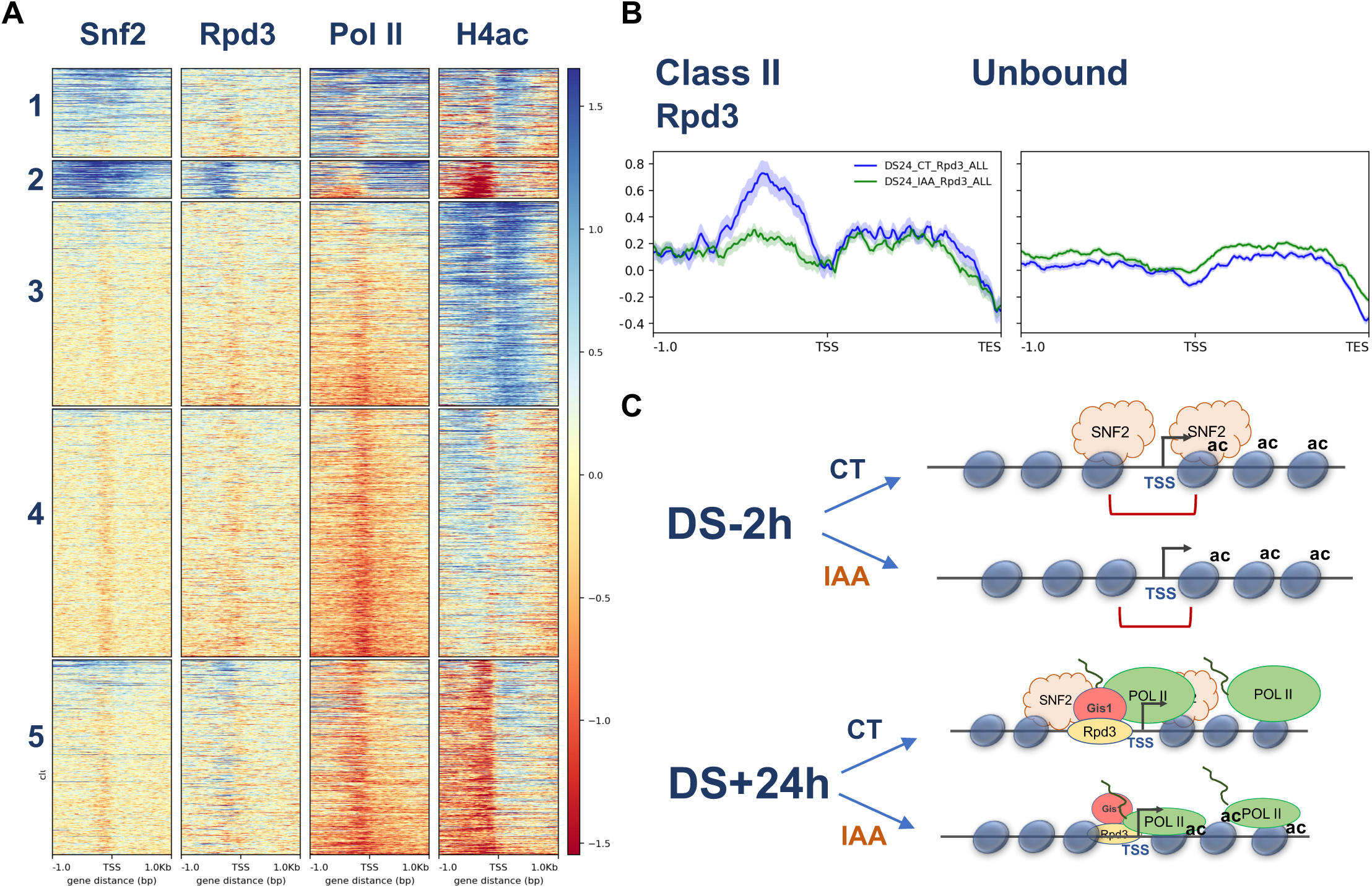
Snf2 facilitates Rpd3 binding to a subset of post-diauxic shift genes. **A)** Heatmap showing K-means cluster analysis of Snf2, Rpd3, Pol II and H4ac ChIP-seq data genome-wide at DS+24h. **B)** Meta-analyses showing Rpd3 binding at DS+24h at Class II (left) and induced unbound genes (right) in control and Snf2-depleted cells **C)** Model showing that Snf2 remodels chromatin at Class II genes at DS-2h (top) to facilitate Gis1 binding, Rpd3 binding, H4ac hypoacetylation and transcription activation after the diauxic shift (bottom).

## Discussion

The decision by a cell to reversibly exit the mitotic cell cycle in response to environmental cues is a robust and highly regulated process. While we know that there is a dramatic global downregulation of transcription as budding yeast cells enter quiescence (McKnight et al., 2015), how the genes that must be activated at various points during this process are turned on is still not well understood. We have identified the SWI/SNF chromatin remodeling complex as an integral player in the activation of Q-specific genes at multiple stages of the Q entry process.

While the SWI/SNF complex was previously implicated in Q cell formation (Li et al., 2015) and stationary phase viability (Reimand et al., 2012), how or when during the 7-day process the activity of SWI/SNF was required was not known. In the waning hours before the DS, cells downregulate genes involved in glycolysis in preparation for the switch to gluconeogenesis and respiration (Brauer et al., 2005; Radonjic et al., 2005; Zampar et al., 2013). During this time, cells also activate genes that promote the cell wall reorganization and fortification required for Q cells to become stress-resistant (Li et al., 2015; Li et al., 2013; Shimoi et al., 1998). We find that Snf2 binds and activates genes required for glycolysis, gluconeogenesis and cell wall reorganization at DS-2h, implying that SWI/SNF is involved in mediating the transition from fermentation to respiration. SWI/SNF has been shown to play important roles during this transition through its effect on splicing and coenzyme Q6 levels (Awad et al., 2017). Our data demonstrate that SWI/SNF directly promotes the transition through the diauxic shift by activating transcription of required genes. This finding is consistent with known roles for SWI/SNF in regulating transcription of stress response genes and metabolic transitions in both yeast and mammals (Dutta et al., 2014; Erkina et al., 2008; Erkina et al., 2010; Neigeborn and Carlson, 1984; Shivaswamy and Iyer, 2008; Wang et al., 2018).

In addition to targeting and regulating genes during the diauxic shift, Snf2 binds many other genes at DS-2h that are regulated at later time points. Of particular interest are the genes bound by Snf2 at DS-2h that are activated after the diauxic shift (post-diauxic shift genes). In yeast, the Rim15 kinase operates under the Ras-cAMP pathway downstream of PKA (Reinders et al., 1998), and upstream of the transcription factors Msn2/4 and Gis1 (Cameroni et al., 2004). We observe SWI/SNF-dependent activation of Msn2 targets at DS-2h. Additionally, Snf2 binds to Gis1 target genes at DS-2h and facilitates both Gis1 binding to their promoters, and transcription later on. Together, these results suggest that SWI/SNF responds to signals from these kinase cascades to activate target gene transcription at the appropriate times. These results are reminiscent of the role of SWI/SNF in responding to signaling cascades to facilitate differentiation. For instance, the Baf60 component of the mammalian SWI/SNF complex binds to myogenic gene promoters prior to the onset of differentiation signaling for chromatin to be remodeled by the catalytic subunits after signaling has been initiated (Forcales, 2012; Harada et al., 2017; Simone et al., 2004). Additionally, mammalian Brg1 is phosphorylated by PKC β1 and dephosphorylated by calcineurin to regulate skeletal muscle differentiation (Nasipak et al., 2015). Overall, our results support a conserved and integral role for SWI/SNF in mediating the transcriptional output of signaling cascades to facilitate cellular proliferation decisions.

As described earlier, quiescence entry is characterized by a massive downregulation in transcription mediated by global histone deacetylation via the histone deacetylase Rpd3 (McKnight et al., 2015). Additionally, cells depleted of glucose experience decreased levels of acetyl-coA and more compact chromatin, creating an environment that is relatively incompatible with transcription (Cai et al., 2011; McKnight et al., 2015; Piñon, 1978; Rutledge et al., 2015; Shi and Tu, 2013). Global histone deacetylation has also been observed in glucose-depleted cells in both mammals and yeast as a method for buffering decreasing intracellular pH (McBrian et al., 2013; Peng et al., 2008). The cytosolic pH of yeast decreases from neutral to 5.5 as yeast enter stationary phase, and cytosolic pH has been linked to the regulation of cellular growth by signal transduction pathways (Orij et al., 2012). Together, these observations support a hypothesis that chromatin acetylation and specific gene regulation mediated by the activity of HATs and HDACs are linked to the sensing of intracellular pH during nutrient changes (Kurdistani, 2014; Kurdistani et al., 2004). In support of this hypothesis, a set of post-diauxic shift genes dubbed histone hypo-acetylation-activated genes (HHAAG) (Mehrotra et al., 2014), some of which are regulated by Snf2, are activated by hypoacetylation upon glucose depletion. Our observation that Snf2 binds at DS-2h to mediate activator binding and the binding and/or activity of Rpd3 at specific post-diauxic shift genes later on, further supports this hypothesis, and provides a mechanism by which gene-specific activation can occur under conditions of global histone deacetylation and low levels of acetyl-CoA. While promoter-bound Rpd3 is most commonly associated with transcriptional repression, there are also instances where it has been shown to bind to active gene promoters or even to mediate activation (Alejandro-Osorio et al., 2009; Borecka-Melkusova et al., 2008; Charles et al., 2011; De Nadal et al., 2004; Kurdistani et al., 2002; Sertil et al., 2007; Sharma et al., 2007). The majority of these genes are involved in the response to stress or nutritional changes, similar to the Class II genes that we have identified. While the increase in H4 acetylation is predominantly in the coding region, the reduced level of Rpd3 at the promoters of Class II genes in Snf2-depleted cells corresponds well to this increase, suggesting that promoter-bound Rpd3 is somehow involved in mediating deacetylation of H4 in coding regions.

Overall, our results implicate Snf2 as a key regulator of transcription activation during the quiescence entry process. The binding patterns of Snf2 as well as the transcriptional and chromatin changes that occur upon Snf2-depletion suggest that Snf2 plays a major role in setting up chromatin architecture early in the quiescence entry process to facilitate events that occur throughout. The coordination between Snf2, Gis1, Rpd3 and H4 acetylation in the coding regions of post-diauxic shift genes, illuminates a previously unknown unique mechanism by which Snf2 responds to the signaling cascades involved in regulating cellular proliferation to activate transcription of specific genes in a relatively repressive environment.

## Acknowledgments

We are very grateful to the members of the Tsukiyama Lab for helpful discussions, especially Sam Cutler, Christine Cucinotta and Jordan Gessaman for help with data analysis. We would also like to acknowledge the Genomics Core at the Fred Hutchinson Cancer Research Center, especially Alyssa Dawson, Andy Marty, Rhonda Meredith and Ryan Basom for help in running and analyzing genomic data. We additionally would like to thank Dr. Steven Hahn and Dr. Chhabi Govind for helpful comments on the manuscript, and Dr. Rafal Donczew for discussing data analyses. This research was done with funding provided by the National Institutes of Health, National Institute of General Medical Sciences (R01-GM058465 and R01-GM111428 to T.T. and F32-GM120955 to M.M.S.

## Author Contributions

All of the authors aided in writing and editing the manuscript. All of the authors discussed experimental design. All of the authors participated in strain construction. Quiescence entry experiments were performed by M.M.S. and K.C.A.B. FACS analyses were performed by K.C.A.B. The chromatin immunoprecipitation experiments for Snf2 in Figure 3 were performed by K.C.A.B. The remaining chromatin immunoprecipitation experiments were performed by M.M.S. 4-TU RNA-seq experiments were performed by M.M.S. Data analysis was performed by M.M.S and K.C.A.B. T.T supervised K.C.A.B. and M.M.S.

## Declaration of Competing Interests

The authors declare no competing interests.

## Figure Legends

**Figure S1.**
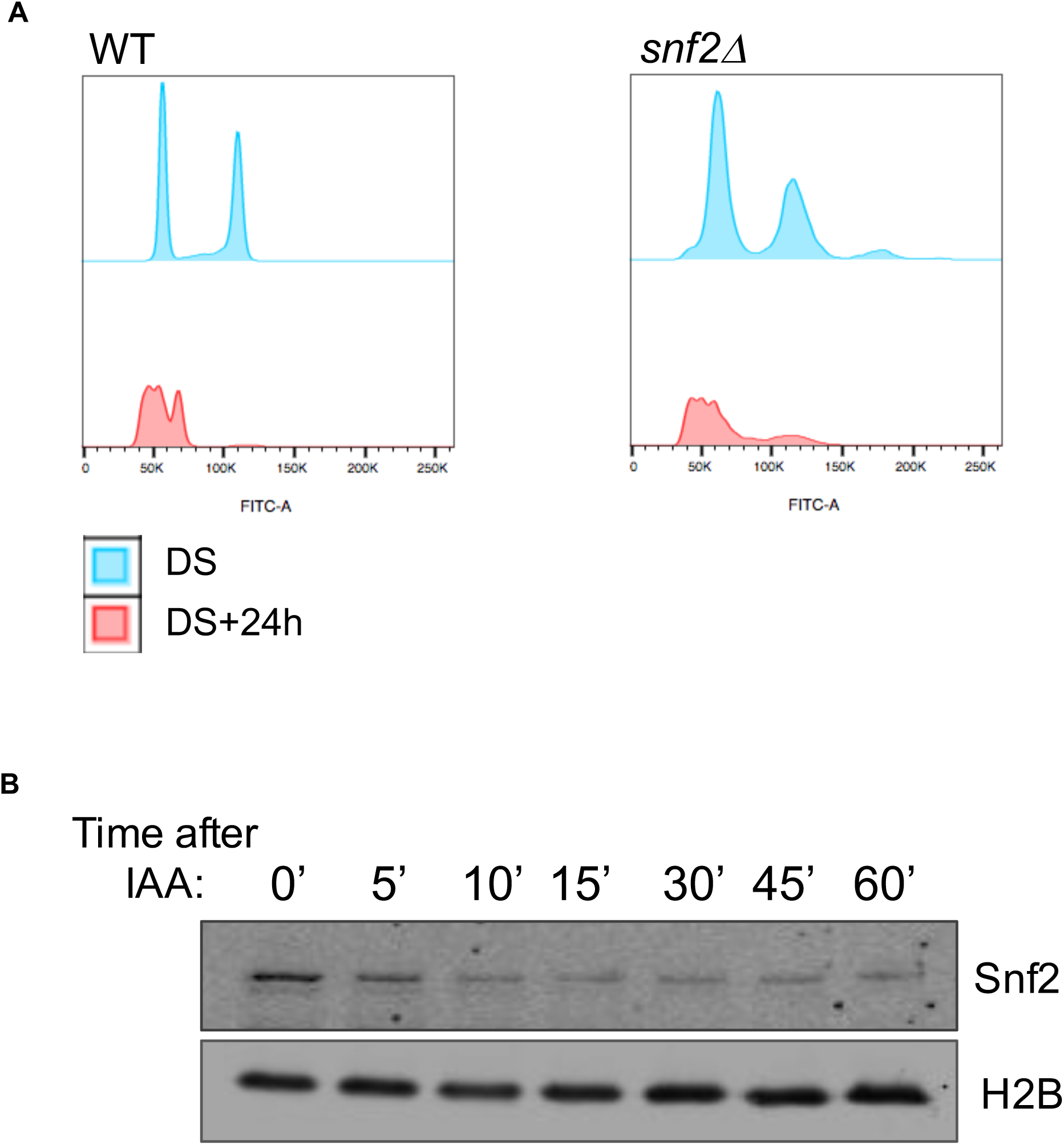
Snf2 is required for Q entry. Cell cycle profiles measured by FACS of WT (left) and *snf2*Δ cells (right) at the DS (blue) and DS+24h (orange) indicating that both cell types gradually arrest in G1 after the DS.

**Figure S2.**
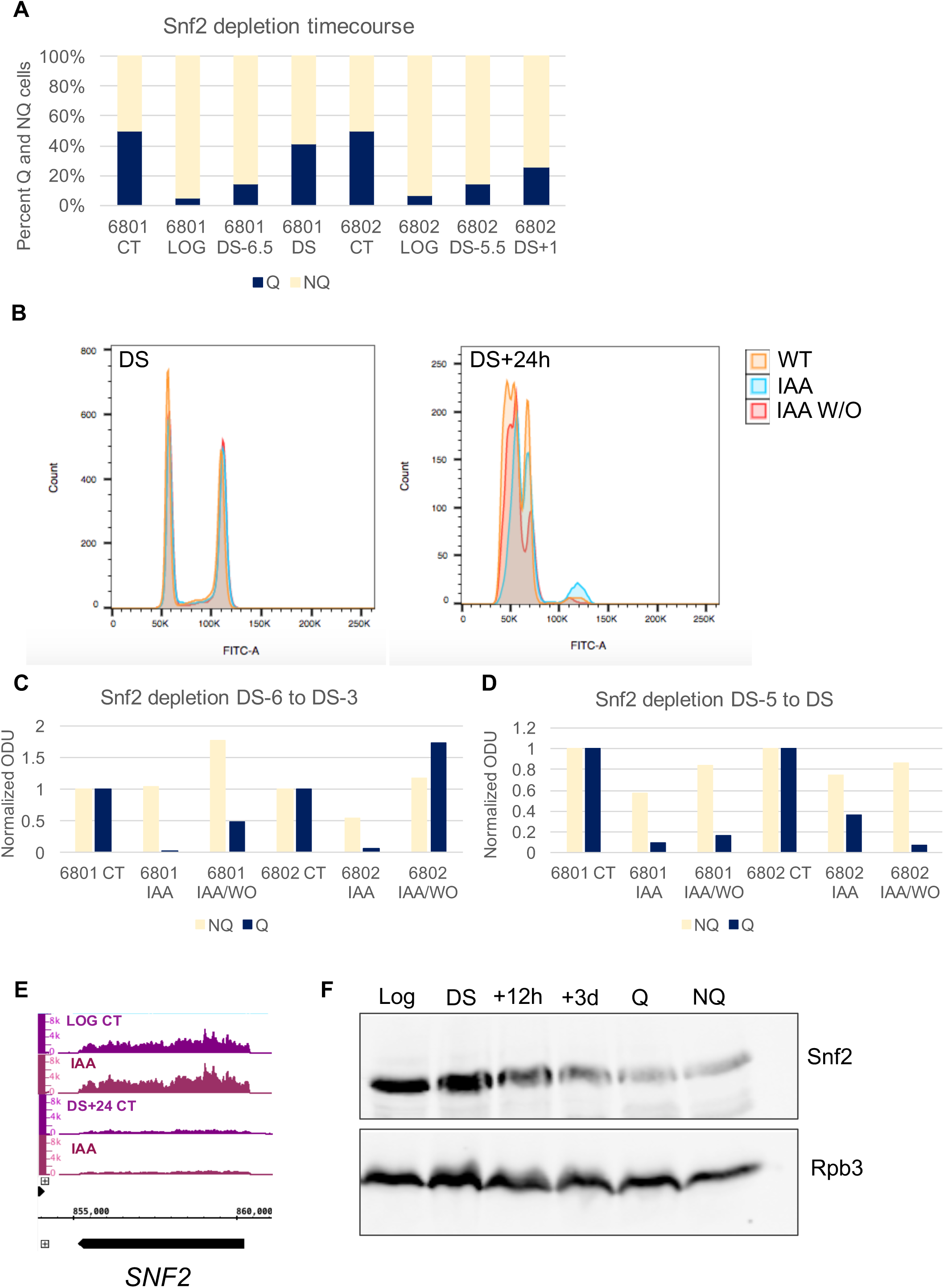
The requirement for Snf2 during Q entry is strongest just prior to the DS. **A)** Percent Q and NQ cells produced by cultures treated with auxin at the indicated times. Auxin remained in the media until the cells were harvested on Day 7 after inoculation. **B)** FACS analysis indicating that cells from which the IAA was washed out have a similar cell cycle profile to that of the untreated cells. At DS+24h, the wash out cells also exhibit the leftward shift in signal indicative of Q cell walls. **C and D)** Normalized OD units of NQ and Q cells purified for the experiments in Figure 2B and C. **E)** 4TU RNA-seq data at the *SNF2* locus in LOG and DS+24h cells untreated (CT) or treated with auxin (IAA). **F)** Western blot of FLAG-tagged Snf2 from samples harvested at the indicated times. Antibody to Rpb3 was used as a control.

**Figure S3.**
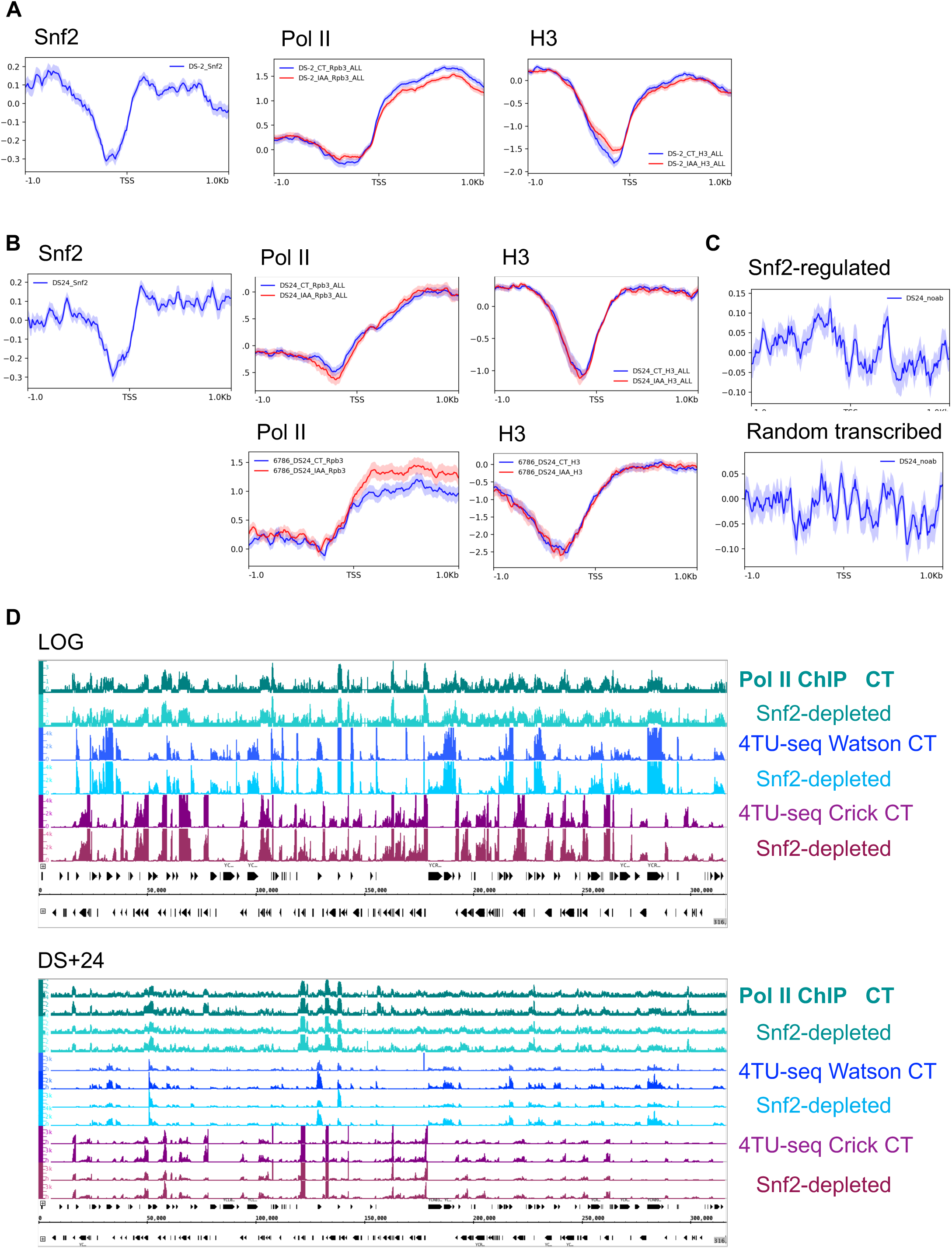
Snf2 binds and regulates genes required for passage through the diauxic shift. **A)** Meta-analyses showing Snf2 binding, Pol II, and histone H3 binding in CT and Snf2-depleted cells at a set of genes with Snf2 binding of less than log_2_ 0.75 that are transcribed at DS-2h. **B)** Meta-analyses showing Snf2 binding, Pol II, and histone H3 binding at a set of randomly chosen genes with less than log2 0.75 Snf2 binding and Pol II occupancy of greater than log2 0.25. Meta-analyses at Snf2-regulated genes or randomly chosen genes transcribed at DS+24h of ChIP-seq in Snf2-FLAG cells using only the Protein-G beads (no primary antibody) (far right top and bottom). **C)** Screen shots showing the correlation between Pol II occupancy in CT and Snf2-depleted cells (green) and strand-specific 4TU-RNA-seq data (blue-Watson strand; burgundy-Crick strand) in log phase cells (top) and at DS+24h (bottom).

**Figure S4.**
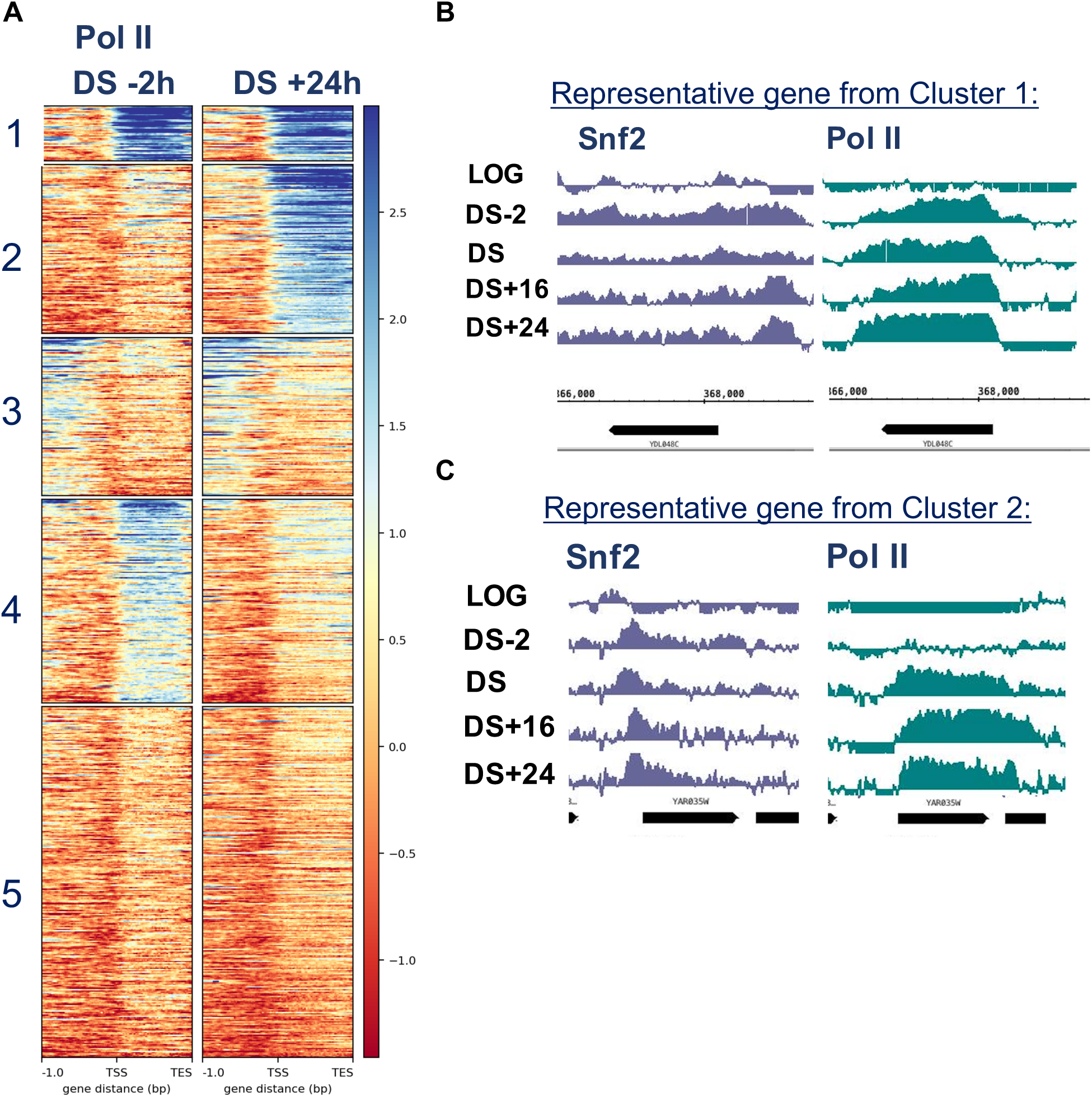
Genes bound by Snf2 at DS-2 exhibit distinct transcription patterns. **A)** Heatmap showing K-means clustering of Pol II (Rpb3) ChIP-seq data at DS-2h and DS+24h at genes bound by Snf2 at DS-2. **B)** Screen shot showing Snf2 and Pol II occupancy at a gene in Cluster 1. **C)** Snf2 and Pol II occupancy at a gene from Cluster 2.

**Figure S5.**
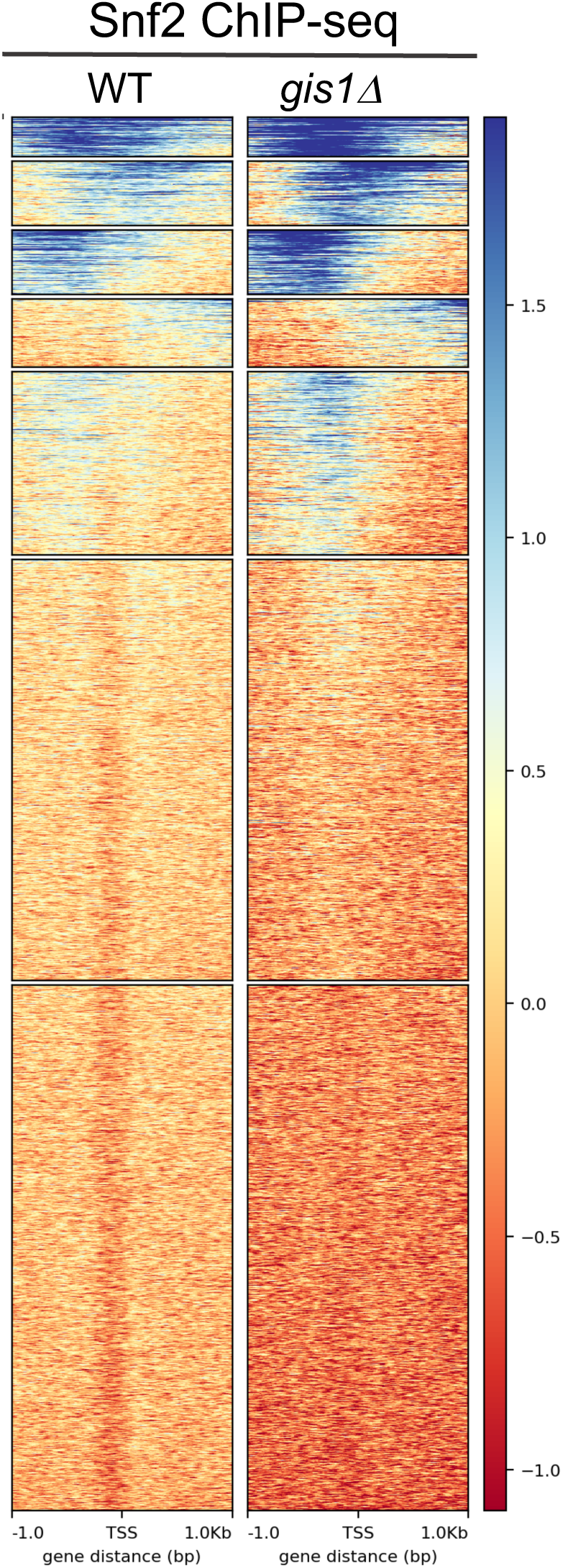
Snf2 facilitates activator binding at post-diauxic shift genes. Heatmap of K-means clustered Snf2 ChIP-seq data in *gis1*Δ cells at DS+24h.

**Figure S6.**
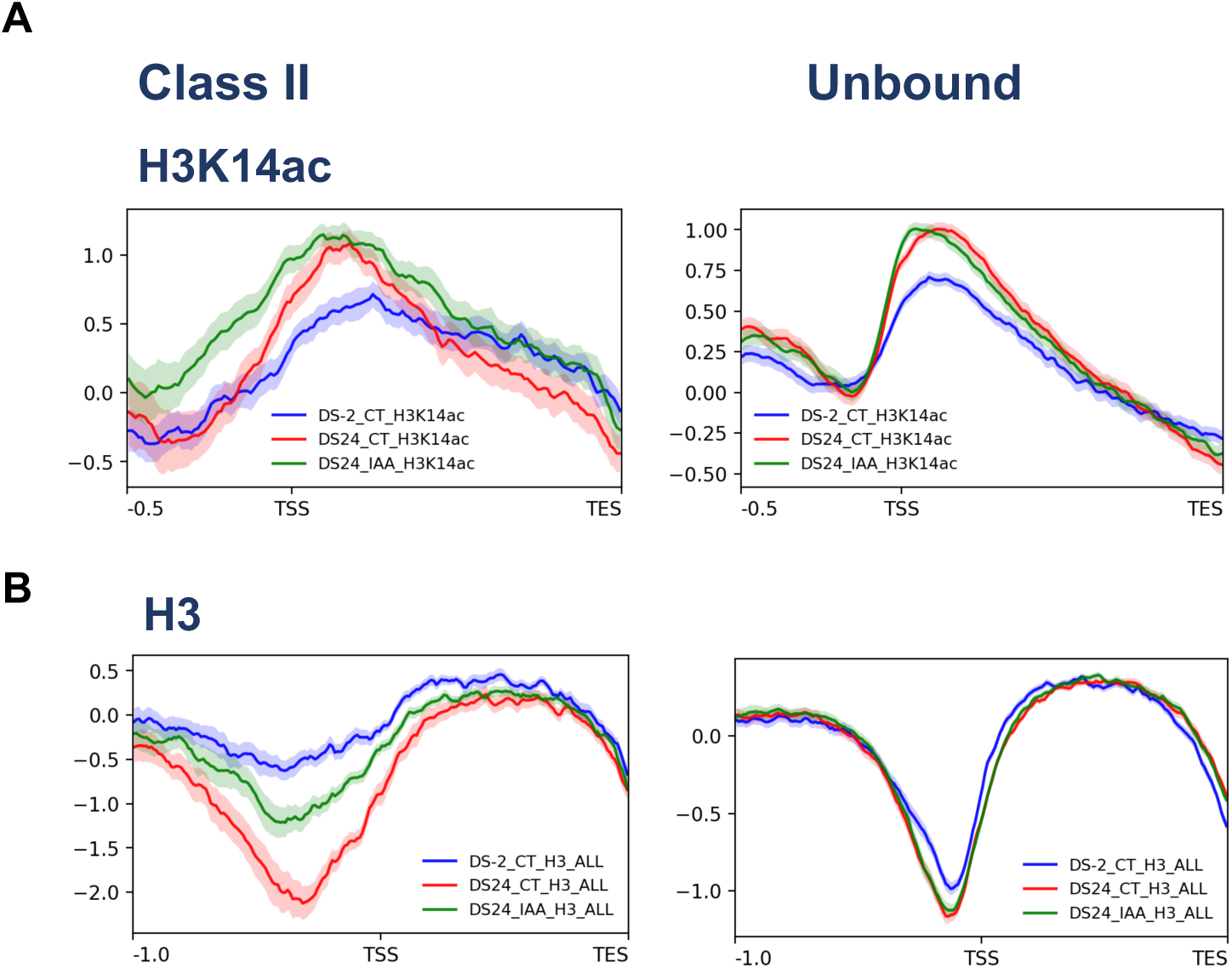
Snf2 promotes histone hypoacetylation in the coding regions of post-diauxic shift genes. **A)** Meta-analyses showing H3K14ac ChIP-seq data at DS+24h at Class II (left) and Snf2-unbound genes that are also transcriptionally induced after the DS (right). **B)** The same for total H3 ChIP-seq data.

## Methods

### Yeast strains and growth conditions

Prototrophic WT RAD5+ W303 strains derived from parent BY6500 (Miles et al., 2013) were used for all of the experiments. The following conditions were used for growing cells for chromatin preparations and Q cell assays. Cells from overnight starter cultures were inoculated at 0.0001 OD_600_ into 40 ml of YPD in a 125 ml Erlenmeyer flask and grown at 30° C with shaking at about 180 rpm. The diauxic shift was monitored using glucose strips (Precision Laboratories). For auxin treatments, 3.6 mg of 3-Indoleacetic acid (Sigma I3750) was directly added to the 40 ml culture since even small amounts of solvents such as water, ethanol, methanol or DMSO added prior to the DS affected Q yield.

### Quiescence entry assays

Assays to measure quiescence entry were performed as described before (Allen et al., 2006; Li et al., 2009; Li et al., 2013; McKnight et al., 2015; Spain et al., 2018). Essentially, cells grown as described above were grown for 7 days. 15 ml of each 40 ml culture was purified over a 12.5 ml Percoll density gradient in a 30 ml glass centrifuge tube. The cells were washed with MilliQ water and the ODU/uL was calculated for both Q and NQ cells.

### ChIP-seq experiments and analysis

ChIP-seq experiments were performed as described previously (McKnight et al., 2015). Antibodies used were anti-H3 (abcam 1791), anti-Rpb3 (Biolegend 665003), anti-Myc (Covance 9E10), anti-H4ac (Upstate 06-866), and anti-FLAG (Sigma F3165). *S. pombe* chromatin was added to 1.25 µg of the chromatin preparations prior to the IP as a spike-in control. Libraries were prepared using the Ovation Ultralow Kit (NuGen). Single end sequencing was done on an Illumina HiSeq 2500 on rapid run mode at the Fred Hutchinson Cancer Research Center Genomics Core. Reads were aligned to the sacCer3 reference genome using bowtie2 version 2.2.3. Coverage files were generated, and immunoprecipitated data were divided by input data using deepTools v.2.0 (Ramírez et al., 2016). Metaplots and heatmaps were also generated using deepTools. Box plots were generated using the ggplot2 function of R.

#### 4TU RNA-seq

Cells were grown with 4TU, RNA was isolated, biotinylated and purified as described previously (Warfield et al., 2017). We began the RNA preparations with a mixture of 1:10 labeled *s. pombe* cells to unlabeled or labeled s. cerevisiae cells. cDNA libraries from 5 ng of cDNA were prepared using the Ovation Solo RNA-seq Kit (NuGen: 0407-32), as per the manufacturer instructions, except that 0.5 µg of actinomycin D was added instead of H2O during Step C “DNase Treatment and Primer Annealing-Total RNA Input”. Paired end sequencing was done on an Illumina HiSeq 2500 at the Fred Hutchinson Cancer Research Center Genomics Core. The initial ratio of *s. pombe* to *s.cerevisiae* in the input cDNA as assessed by RT-PCR was used to normalize the reads. Reads were aligned to the sacCer3 reference genome using HISAT2 on the public server of Galaxy (*usegalaxy.org* (Afgan et al., 2016)). Reads from unlabeled samples were substracted from that of labeled samples using the BamCompare function of deepTools in Galaxy with RPKM normalization, with the scaling factor determined from the input normalization step. Lastly, snf2-depleted samples were normalized to control samples by using the BigWigCompare function in deepTools with the scaling factor determined by normalizing the snf2-depleted versus control percent reads ratio of *s. cerevisiae* to *s. pombe* in the immunoprecipitated samples.

## References

Abdulrehman, D., Monteiro, P.T., Teixeira, M.C., Mira, N.P., Lourenço, A.B., dos Santos, S.C., Cabrito, T.R., Francisco, A.P., Madeira, S.C., Aires, R.S., et al. (2011). YEASTRACT: providing a programmatic access to curated transcriptional regulatory associations in Saccharomyces cerevisiae through a web services interface. Nucleic Acids Res 39, D136–140.

Abrams, E., Neigeborn, L., and Carlson, M. (1986). Molecular analysis of SNF2 and SNF5, genes required for expression of glucose-repressible genes in Saccharomyces cerevisiae. Mol Cell Biol 6, 3643–3651.

Afgan, E., Baker, D., van den Beek, M., Blankenberg, D., Bouvier, D., Čech, M., Chilton, J., Clements, D., Coraor, N., Eberhard, C., et al. (2016). The Galaxy platform for accessible, reproducible and collaborative biomedical analyses: 2016 update. Nucleic Acids Res 44, W3–W10.

Alejandro-Osorio, A.L., Huebert, D.J., Porcaro, D.T., Sonntag, M.E., Nillasithanukroh, S., Will, J.L., and Gasch, A.P. (2009). The histone deacetylase Rpd3p is required for transient changes in genomic expression in response to stress. Genome Biol 10, R57.

Allen, C., Büttner, S., Aragon, A.D., Thomas, J.A., Meirelles, O., Jaetao, J.E., Benn, D., Ruby, S.W., Veenhuis, M., Madeo, F., et al. (2006). Isolation of quiescent and nonquiescent cells from yeast stationary-phase cultures. J Cell Biol 174, 89–100.

Amrani, Y.M., Gill, J., Matevossian, A., Alonzo, E.S., Yang, C., Shieh, J.H., Moore, M.A., Park, C.Y., Sant’Angelo, D.B., and Denzin, L.K. (2011). The Paf oncogene is essential for hematopoietic stem cell function and development. J Exp Med 208, 1757–1765.

Aragon, A.D., Rodriguez, A.L., Meirelles, O., Roy, S., Davidson, G.S., Tapia, P.H., Allen, C., Joe, R., Benn, D., and Werner-Washburne, M. (2008). Characterization of differentiated quiescent and nonquiescent cells in yeast stationary-phase cultures. Mol Biol Cell 19, 1271–1280.

Awad, A.M., Venkataramanan, S., Nag, A., Galivanche, A.R., Bradley, M.C., Neves, L.T., Douglass, S., Clarke, C.F., and Johnson, T.L. (2017). Chromatin-remodeling SWI/SNF complex regulates coenzyme Q. J Biol Chem 292, 14851–14866.

Bischoff, R. (1990). Cell cycle commitment of rat muscle satellite cells. J Cell Biol 111, 201–207.

Bontron, S., Jaquenoud, M., Vaga, S., Talarek, N., Bodenmiller, B., Aebersold, R., and De Virgilio, C. (2013). Yeast endosulfines control entry into quiescence and chronological life span by inhibiting protein phosphatase 2A. Cell Rep 3, 16–22.

Borecka-Melkusova, S., Kozovska, Z., Hikkel, I., Dzugasova, V., and Subik, J. (2008). RPD3 and ROM2 are required for multidrug resistance in Saccharomyces cerevisiae. FEMS Yeast Res 8, 414–424.

Brauer, M.J., Saldanha, A.J., Dolinski, K., and Botstein, D. (2005). Homeostatic adjustment and metabolic remodeling in glucose-limited yeast cultures. Mol Biol Cell 16, 2503–2517.

Burns, L.G., and Peterson, C.L. (1997). The yeast SWI-SNF complex facilitates binding of a transcriptional activator to nucleosomal sites in vivo. Mol Cell Biol 17, 4811–4819.

Cai, L., Sutter, B.M., Li, B., and Tu, B.P. (2011). Acetyl-CoA induces cell growth and proliferation by promoting the acetylation of histones at growth genes. Mol Cell 42, 426–437.

Cameroni, E., Hulo, N., Roosen, J., Winderickx, J., and De Virgilio, C. (2004). The novel yeast PAS kinase Rim 15 orchestrates G0-associated antioxidant defense mechanisms. Cell Cycle 3, 462–468.

Chandy, M., Gutierrez, J.L., Prochasson, P., and Workman, J.L. (2006). SWI/SNF displaces SAGA-acetylated nucleosomes. Eukaryot Cell 5, 1738–1747.

Charles, G.M., Chen, C., Shih, S.C., Collins, S.R., Beltrao, P., Zhang, X., Sharma, T., Tan, S., Burlingame, A.L., Krogan, N.J., et al. (2011). Site-specific acetylation mark on an essential chromatin-remodeling complex promotes resistance to replication stress. Proceedings of the National Academy of Sciences of the United States of America 108, 10620–10625.

Cheung, T.H., and Rando, T.A. (2013). Molecular regulation of stem cell quiescence. Nat Rev Mol Cell Biol 14, 329–340.

Coller, H.A., Sang, L., and Roberts, J.M. (2006). A new description of cellular quiescence. PLoS Biol 4, e83.

Cosma, M.P., Tanaka, T., and Nasmyth, K. (1999). Ordered recruitment of transcription and chromatin remodeling factors to a cell cycle-and developmentally regulated promoter. Cell 97, 299–311.

Côté, J., Quinn, J., Workman, J.L., and Peterson, C.L. (1994). Stimulation of GAL4 derivative binding to nucleosomal DNA by the yeast SWI/SNF complex. Science 265, 53–60.

De Nadal, E., Zapater, M., Alepuz, P.M., Sumoy, L., Mas, G., and Posas, F. (2004). The MAPK Hog1 recruits Rpd3 histone deacetylase to activate osmoresponsive genes. Nature 427, 370–374.

De Virgilio, C. (2012). The essence of yeast quiescence. FEMS Microbiol Rev 36, 306–339.

De Virgilio, C., and Loewith, R. (2006). The TOR signalling network from yeast to man. Int J Biochem Cell Biol 38, 1476–1481.

DeRisi, J.L., Iyer, V.R., and Brown, P.O. (1997). Exploring the metabolic and genetic control of gene expression on a genomic scale. Science 278, 680–686.

Dhawan, J., and Laxman, S. (2015). Decoding the stem cell quiescence cycle--lessons from yeast for regenerative biology. J Cell Sci 128, 4467–4474.

Drouin, S., Laramée, L., Jacques, P., Forest, A., Bergeron, M., and Robert, F. (2010). DSIF and RNA polymerase II CTD phosphorylation coordinate the recruitment of Rpd3S to actively transcribed genes. PLoS Genet 6, e1001173.

Duffy, E.E., Rutenberg-Schoenberg, M., Stark, C.D., Kitchen, R.R., Gerstein, M.B., and Simon, M.D. (2015). Tracking Distinct RNA Populations Using Efficient and Reversible Covalent Chemistry. Mol Cell 59, 858–866.

Duffy, E.E., and Simon, M.D. (2016). Enriching s. Curr Protoc Chem Biol 8, 234–250.

Dutta, A., Gogol, M., Kim, J.H., Smolle, M., Venkatesh, S., Gilmore, J., Florens, L., Washburn, M.P., and Workman, J.L. (2014). Swi/Snf dynamics on stress-responsive genes is governed by competitive bromodomain interactions. Genes Dev 28, 2314–2330.

Erkina, T.Y., Tschetter, P.A., and Erkine, A.M. (2008). Different requirements of the SWI/SNF complex for robust nucleosome displacement at promoters of heat shock factor and Msn2-and Msn4-regulated heat shock genes. Mol Cell Biol 28, 1207–1217.

Erkina, T.Y., Zou, Y., Freeling, S., Vorobyev, V.I., and Erkine, A.M. (2010). Functional interplay between chromatin remodeling complexes RSC, SWI/SNF and ISWI in regulation of yeast heat shock genes. Nucleic Acids Res 38, 1441–1449.

Fabrizio, P., Pozza, F., Pletcher, S.D., Gendron, C.M., and Longo, V.D. (2001). Regulation of longevity and stress resistance by Sch9 in yeast. Science 292, 288–290.

Fausto, N. (2004). Liver regeneration and repair: hepatocytes, progenitor cells, and stem cells. Hepatology 39, 1477–1487.

Forcales, S.V. (2012). The BAF60c-MyoD complex poises chromatin for rapid transcription. Bioarchitecture 2, 104–109.

François, J., and Parrou, J.L. (2001). Reserve carbohydrates metabolism in the yeast Saccharomyces cerevisiae. FEMS Microbiol Rev 25, 125–145.

Galdieri, L., Mehrotra, S., Yu, S., and Vancura, A. (2010). Transcriptional regulation in yeast during diauxic shift and stationary phase. OMICS 14, 629–638.

Gasch, A.P., Spellman, P.T., Kao, C.M., Carmel-Harel, O., Eisen, M.B., Storz, G., Botstein, D., and Brown, P.O. (2000). Genomic expression programs in the response of yeast cells to environmental changes. Mol Biol Cell 11, 4241–4257.

Geng, F., and Laurent, B.C. (2004). Roles of SWI/SNF and HATs throughout the dynamic transcription of a yeast glucose-repressible gene. EMBO J 23, 127–137.

Govind, C.K., Qiu, H., Ginsburg, D.S., Ruan, C., Hofmeyer, K., Hu, C., Swaminathan, V., Workman, J.L., Li, B., and Hinnebusch, A.G. (2010). Phosphorylated Pol II CTD recruits multiple HDACs, including Rpd3C(S), for methylation-dependent deacetylation of ORF nucleosomes. Mol Cell 39, 234–246.

Gray, J.V., Petsko, G.A., Johnston, G.C., Ringe, D., Singer, R.A., and Werner-Washburne, M. (2004). “Sleeping beauty”: quiescence in Saccharomyces cerevisiae. Microbiol Mol Biol Rev 68, 187–206.

Harada, A., Ohkawa, Y., and Imbalzano, A.N. (2017). Temporal regulation of chromatin during myoblast differentiation. Semin Cell Dev Biol 72, 77–86.

Hassan, A.H., Awad, S., and Prochasson, P. (2006). The Swi2/Snf2 bromodomain is required for the displacement of SAGA and the octamer transfer of SAGA-acetylated nucleosomes. J Biol Chem 281, 18126–18134.

Hassan, A.H., Prochasson, P., Neely, K.E., Galasinski, S.C., Chandy, M., Carrozza, M.J., and Workman, J.L. (2002). Function and selectivity of bromodomains in anchoring chromatin-modifying complexes to promoter nucleosomes. Cell 111, 369–379.

Imbalzano, A.N., Kwon, H., Green, M.R., and Kingston, R.E. (1994). Facilitated binding of TATA-binding protein to nucleosomal DNA. Nature 370, 481–485.

Keogh, M.C., Kurdistani, S.K., Morris, S.A., Ahn, S.H., Podolny, V., Collins, S.R., Schuldiner, M., Chin, K., Punna, T., Thompson, N.J., et al. (2005). Cotranscriptional set2 methylation of histone H3 lysine 36 recruits a repressive Rpd3 complex. Cell 123, 593–605.

Kurdistani, S.K. (2014). Chromatin: a capacitor of acetate for integrated regulation of gene expression and cell physiology. Curr Opin Genet Dev 26, 53–58.

Kurdistani, S.K., Robyr, D., Tavazoie, S., and Grunstein, M. (2002). Genome-wide binding map of the histone deacetylase Rpd3 in yeast. Nat Genet 31, 248–254.

Kurdistani, S.K., Tavazoie, S., and Grunstein, M. (2004). Mapping global histone acetylation patterns to gene expression. Cell 117, 721–733.

Kwon, H., Imbalzano, A.N., Khavari, P.A., Kingston, R.E., and Green, M.R. (1994). Nucleosome disruption and enhancement of activator binding by a human SW1/SNF complex. Nature 370, 477–481.

Li, B., Gogol, M., Carey, M., Pattenden, S.G., Seidel, C., and Workman, J.L. (2007). Infrequently transcribed long genes depend on the Set2/Rpd3S pathway for accurate transcription. Genes Dev 21, 1422–1430.

Li, L., Lu, Y., Qin, L.X., Bar-Joseph, Z., Werner-Washburne, M., and Breeden, L.L. (2009). Budding yeast SSD1-V regulates transcript levels of many longevity genes and extends chronological life span in purified quiescent cells. Mol Biol Cell 20, 3851–3864.

Li, L., Miles, S., and Breeden, L.L. (2015). A Genetic Screen for Saccharomyces cerevisiae Mutants That Fail to Enter Quiescence. G3 (Bethesda) 5, 1783–1795.

Li, L., Miles, S., Melville, Z., Prasad, A., Bradley, G., and Breeden, L.L. (2013). Key events during the transition from rapid growth to quiescence in budding yeast require posttranscriptional regulators. Mol Biol Cell 24, 3697–3709.

Lillie, S.H., and Pringle, J.R. (1980). Reserve carbohydrate metabolism in Saccharomyces cerevisiae: responses to nutrient limitation. J Bacteriol 143, 1384–1394.

Longo, V.D., and Fabrizio, P. (2012). Chronological aging in Saccharomyces cerevisiae. Subcell Biochem 57, 101–121.

Martínez-Pastor, M.T., Marchler, G., Schüller, C., Marchler-Bauer, A., Ruis, H., and Estruch, F. (1996). The Saccharomyces cerevisiae zinc finger proteins Msn2p and Msn4p are required for transcriptional induction through the stress response element (STRE). EMBO J 15, 2227–2235.

McBrian, M.A., Behbahan, I.S., Ferrari, R., Su, T., Huang, T.W., Li, K., Hong, C.S., Christofk, H.R., Vogelauer, M., Seligson, D.B., et al. (2013). Histone acetylation regulates intracellular pH. Mol Cell 49, 310–321.

McKnight, J.N., Boerma, J.W., Breeden, L.L., and Tsukiyama, T. (2015). Global Promoter Targeting of a Conserved Lysine Deacetylase for Transcriptional Shutoff during Quiescence Entry. Mol Cell 59, 732–743.

Mehrotra, S., Galdieri, L., Zhang, T., Zhang, M., Pemberton, L.F., and Vancura, A. (2014). Histone hypoacetylation-activated genes are repressed by acetyl-CoA-and chromatin-mediated mechanism. Biochim Biophys Acta 1839, 751–763.

Miles, S., and Breeden, L. (2016). A common strategy for initiating the transition from proliferation to quiescence. Curr Genet.

Miles, S., and Breeden, L. (2017). A common strategy for initiating the transition from proliferation to quiescence. Curr Genet 63, 179–186.

Miles, S., Li, L., Davison, J., and Breeden, L.L. (2013). Xbp1 directs global repression of budding yeast transcription during the transition to quiescence and is important for the longevity and reversibility of the quiescent state. PLoS Genet 9, e1003854.

Mitra, D., Parnell, E.J., Landon, J.W., Yu, Y., and Stillman, D.J. (2006). SWI/SNF binding to the HO promoter requires histone acetylation and stimulates TATA-binding protein recruitment. Mol Cell Biol 26, 4095–4110.

Monteiro, P.T., Mendes, N.D., Teixeira, M.C., ďOrey, S., Tenreiro, S., Mira, N.P., Pais, H., Francisco, A.P., Carvalho, A.M., Lourenço, A.B., et al. (2008). YEASTRACT-DISCOVERER: new tools to improve the analysis of transcriptional regulatory associations in Saccharomyces cerevisiae. Nucleic Acids Res 36, D132–136.

Nasipak, B.T., Padilla-Benavides, T., Green, K.M., Leszyk, J.D., Mao, W., Konda, S., Sif, S., Shaffer, S.A., Ohkawa, Y., and Imbalzano, A.N. (2015). Opposing calcium-dependent signalling pathways control skeletal muscle differentiation by regulating a chromatin remodelling enzyme. Nat Commun 6, 7441.

Neigeborn, L., and Carlson, M. (1984). Genes affecting the regulation of SUC2 gene expression by glucose repression in Saccharomyces cerevisiae. Genetics 108, 845–858.

Nishimura, K., Fukagawa, T., Takisawa, H., Kakimoto, T., and Kanemaki, M. (2009). An auxin-based degron system for the rapid depletion of proteins in nonplant cells. Nat Methods 6, 917–922.

Orij, R., Urbanus, M.L., Vizeacoumar, F.J., Giaever, G., Boone, C., Nislow, C., Brul, S., and Smits, G.J. (2012). Genome-wide analysis of intracellular pH reveals quantitative control of cell division rate by pH(c) in Saccharomyces cerevisiae. Genome Biol 13, R80.

Parrou, J.L., Enjalbert, B., and François, J. (1999). STRE-and cAMP-independent transcriptional induction of Saccharomyces cerevisiae GSY2 encoding glycogen synthase during diauxic growth on glucose. Yeast 15, 1471–1484.

Pedruzzi, I., Bürckert, N., Egger, P., and De Virgilio, C. (2000). Saccharomyces cerevisiae Ras/cAMP pathway controls post-diauxic shift element-dependent transcription through the zinc finger protein Gis1. EMBO J 19, 2569–2579.

Peng, L.H., Yang, X.F., and Fu, N. (2008). [The effects of curcumin on the expression of acetyl-CoA carboxylase of HL-7702 cells]. Zhonghua Gan Zang Bing Za Zhi 16, 948–949.

Peterson, C.L., Dingwall, A., and Scott, M.P. (1994). Five SWI/SNF gene products are components of a large multisubunit complex required for transcriptional enhancement. Proceedings of the National Academy of Sciences of the United States of America 91, 2905–2908.

Piñon, R. (1978). Folded chromosomes in non-cycling yeast cells: evidence for a characteristic g0 form. Chromosoma 67, 263–274.

Qiu, H., Chereji, R.V., Hu, C., Cole, H.A., Rawal, Y., Clark, D.J., and Hinnebusch, A.G. (2016). Genome-wide cooperation by HAT Gcn5, remodeler SWI/SNF, and chaperone Ydj1 in promoter nucleosome eviction and transcriptional activation. Genome Res 26, 211–225.

Radonjic, M., Andrau, J.C., Lijnzaad, P., Kemmeren, P., Kockelkorn, T.T., van Leenen, D., van Berkum, N.L., and Holstege, F.C. (2005). Genome-wide analyses reveal RNA polymerase II located upstream of genes poised for rapid response upon S. cerevisiae stationary phase exit. Mol Cell 18, 171–183.

Ramírez, F., Ryan, D.P., Grüning, B., Bhardwaj, V., Kilpert, F., Richter, A.S., Heyne, S., Dündar, F., and Manke, T. (2016). deepTools2: a next generation web server for deep-sequencing data analysis. Nucleic Acids Res 44, W160–165.

Reimand, J., Aun, A., Vilo, J., Vaquerizas, J.M., Sedman, J., and Luscombe, N.M. (2012). m:Explorer: multinomial regression models reveal positive and negative regulators of longevity in yeast quiescence. Genome Biol 13, R55.

Reinders, A., Bürckert, N., Boller, T., Wiemken, A., and De Virgilio, C. (1998). Saccharomyces cerevisiae cAMP-dependent protein kinase controls entry into stationary phase through the Rim15p protein kinase. Genes Dev 12, 2943–2955.

Reinke, H., Gregory, P.D., and Hörz, W. (2001). A transient histone hyperacetylation signal marks nucleosomes for remodeling at the PHO8 promoter in vivo. Mol Cell 7, 529–538.

Rodgers, J.T., King, K.Y., Brett, J.O., Cromie, M.J., Charville, G.W., Maguire, K.K., Brunson, C., Mastey, N., Liu, L., Tsai, C.R., et al. (2014). mTORC1 controls the adaptive transition of quiescent stem cells from G0 to G(Alert). Nature 510, 393–396.

Rutledge, M.T., Russo, M., Belton, J.M., Dekker, J., and Broach, J.R. (2015). The yeast genome undergoes significant topological reorganization in quiescence. Nucleic Acids Res 43, 8299–8313.

Ryan, M.P., Jones, R., and Morse, R.H. (1998). SWI-SNF complex participation in transcriptional activation at a step subsequent to activator binding. Mol Cell Biol 18, 1774–1782.

Sanz, A.B., García, R., Rodríguez-Peña, J.M., Díez-Muñiz, S., Nombela, C., Peterson, C.L., and Arroyo, J. (2012). Chromatin remodeling by the SWI/SNF complex is essential for transcription mediated by the yeast cell wall integrity MAPK pathway. Mol Biol Cell 23, 2805–2817.

Sanz, A.B., García, R., Rodríguez-Peña, J.M., Nombela, C., and Arroyo, J. (2016). Cooperation between SAGA and SWI/SNF complexes is required for efficient transcriptional responses regulated by the yeast MAPK Slt2. Nucleic Acids Res 44, 7159–7172.

Schaniel, C., Sirabella, D., Qiu, J., Niu, X., Lemischka, I.R., and Moore, K.A. (2011). Wnt-inhibitory factor 1 dysregulation of the bone marrow niche exhausts hematopoietic stem cells. Blood 118, 2420–2429.

Sertil, O., Vemula, A., Salmon, S.L., Morse, R.H., and Lowry, C.V. (2007). Direct role for the Rpd3 complex in transcriptional induction of the anaerobic DAN/TIR genes in yeast. Mol Cell Biol 27, 2037–2047.

Sharma, V.M., Tomar, R.S., Dempsey, A.E., and Reese, J.C. (2007). Histone deacetylases RPD3 and HOS2 regulate the transcriptional activation of DNA damage-inducible genes. Mol Cell Biol 27, 3199–3210.

Shi, L., and Tu, B.P. (2013). Acetyl-CoA induces transcription of the key G1 cyclin CLN3 to promote entry into the cell division cycle in Saccharomyces cerevisiae. Proceedings of the National Academy of Sciences of the United States of America 110, 7318–7323.

Shimoi, H., Kitagaki, H., Ohmori, H., Iimura, Y., and Ito, K. (1998). Sed1p is a major cell wall protein of Saccharomyces cerevisiae in the stationary phase and is involved in lytic enzyme resistance. J Bacteriol 180, 3381–3387.

Shivaswamy, S., and Iyer, V.R. (2008). Stress-dependent dynamics of global chromatin remodeling in yeast: dual role for SWI/SNF in the heat shock stress response. Mol Cell Biol 28, 2221–2234.

Simone, C., Forcales, S.V., Hill, D.A., Imbalzano, A.N., Latella, L., and Puri, P.L. (2004). p38 pathway targets SWI-SNF chromatin-remodeling complex to muscle-specific loci. Nat Genet 36, 738–743.

Spain, M.M., Swygert, S.G., and Tsukiyama, T. (2018). Preparation and Analysis of Saccharomyces cerevisiae Quiescent Cells. Methods Mol Biol 1686, 125–135.

Swinnen, E., Wanke, V., Roosen, J., Smets, B., Dubouloz, F., Pedruzzi, I., Cameroni, E., De Virgilio, C., and Winderickx, J. (2006). Rim15 and the crossroads of nutrient signalling pathways in Saccharomyces cerevisiae. Cell Div 1, 3.

Tatchell, K. (1986). RAS genes and growth control in Saccharomyces cerevisiae. J Bacteriol 166, 364–367.

Teixeira, M.C., Monteiro, P., Jain, P., Tenreiro, S., Fernandes, A.R., Mira, N.P., Alenquer, M., Freitas, A.T., Oliveira, A.L., and Sá-Correia, I. (2006). The YEASTRACT database: a tool for the analysis of transcription regulatory associations in Saccharomyces cerevisiae. Nucleic Acids Res 34, D446–451.

Teixeira, M.C., Monteiro, P.T., Guerreiro, J.F., Gonçalves, J.P., Mira, N.P., dos Santos, S.C., Cabrito, T.R., Palma, M., Costa, C., Francisco, A.P., et al. (2014). The YEASTRACT database: an upgraded information system for the analysis of gene and genomic transcription regulation in Saccharomyces cerevisiae. Nucleic Acids Res 42, D161–166.

Teixeira, M.C., Monteiro, P.T., Palma, M., Costa, C., Godinho, C.P., Pais, P., Cavalheiro, M., Antunes, M., Lemos, A., Pedreira, T., et al. (2018). YEASTRACT: an upgraded database for the analysis of transcription regulatory networks in Saccharomyces cerevisiae. Nucleic Acids Res 46, D348–D353.

Teixeira, M.C., Monteiro, P.T., and Sá-Correia, I. (2016). Predicting Gene and Genomic Regulation in Saccharomyces cerevisiae, using the YEASTRACT Database: A Step-by-Step Guided Analysis. Methods Mol Biol 1361, 391–404.

Thevelein, J.M., and de Winde, J.H. (1999). Novel sensing mechanisms and targets for the cAMP-protein kinase A pathway in the yeast Saccharomyces cerevisiae. Mol Microbiol 33, 904–918.

Valcourt, J.R., Lemons, J.M., Haley, E.M., Kojima, M., Demuren, O.O., and Coller, H.A. (2012). Staying alive: metabolic adaptations to quiescence. Cell Cycle 11, 1680–1696.

Wang, R.R., Pan, R., Zhang, W., Fu, J., Lin, J.D., and Meng, Z.X. (2018). The SWI/SNF chromatin-remodeling factors BAF60a, b, and c in nutrient signaling and metabolic control. Protein Cell 9, 207–215.

Warfield, L., Ramachandran, S., Baptista, T., Devys, D., Tora, L., and Hahn, S. (2017). Transcription of Nearly All Yeast RNA Polymerase II-Transcribed Genes Is Dependent on Transcription Factor TFIID. Mol Cell 68, 118–129.e115.

Werner-Washburne, M., Braun, E., Johnston, G.C., and Singer, R.A. (1993). Stationary phase in the yeast Saccharomyces cerevisiae. Microbiol Rev 57, 383–401.

Yen, K., Vinayachandran, V., Batta, K., Koerber, R.T., and Pugh, B.F. (2012). Genome-wide nucleosome specificity and directionality of chromatin remodelers. Cell 149, 1461–1473.

Yoshinaga, S.K., Peterson, C.L., Herskowitz, I., and Yamamoto, K.R. (1992). Roles of SWI1, SWI2, and SWI3 proteins for transcriptional enhancement by steroid receptors. Science 258, 1598–1604.

Yudkovsky, N., Logie, C., Hahn, S., and Peterson, C.L. (1999). Recruitment of the SWI/SNF chromatin remodeling complex by transcriptional activators. Genes Dev 13, 2369–2374.

Zampar, G.G., Kümmel, A., Ewald, J., Jol, S., Niebel, B., Picotti, P., Aebersold, R., Sauer, U., Zamboni, N., and Heinemann, M. (2013). Temporal system-level organization of the switch from glycolytic to gluconeogenic operation in yeast. Mol Syst Biol 9, 651.

Zhang, N., Wu, J., and Oliver, S.G. (2009). Gis1 is required for transcriptional reprogramming of carbon metabolism and the stress response during transition into stationary phase in yeast. Microbiology 155, 1690–1698.

